# Phylogeography and diversification of the *Pieris napi* species group in the Western Palaearctic

**DOI:** 10.1101/2025.01.31.634921

**Authors:** Javier Sala-Garcia, Maria Vives-Ingla, Vanina Tonzo, Vlad Dincă, Roger Vila, Jofre Carnicer

## Abstract

The butterflies of the *Pieris napi* complex, encompassing diverse taxa across Europe, Asia and North Africa, represent an emergent model system for studying climatic adaptation. In this study, we use genomic data for this group to address phylogenetic relationships, population structure and ecological differentiation in the Western Palearctic. Our results reveal two main groups: the *napi* clade and the *bryoniae-balcana* clade. We clarify the status and show that the distribution of the debated species *P. balcana* extends to south Romania and the Peloponnese. This species is shown to be sister to *P. bryoniae*, which indicates that melanism is an evolutionarily labile trait linked to climatic adaptation. The *P. napi* clade is divided into three components, each predominating in a different south European peninsula, and the boreal *P. n. adalwinda*, which is notably differentiated. The endangered Moroccan taxon *segonzaci* produced conflicting results, probably due to low-quality DNA. Admixture analyses suggest substantial gene flow between taxa in contact areas. Redundancy analyses identify summer temperature and precipitation as key drivers of adaptive genetic variation within the group. This work provides a robust evolutionary framework for future ecological studies of the *Pieris napi* complex, to forecast eco-evolutionary responses and to address conservation priorities in a rapidly changing climate.

## Introduction

European butterflies have become a model taxon for biodiversity and conservation studies, monitoring of insect populations, and global change impact studies (Hanski, 1999; Hanski & Mononen, 2011; Parmesan et al., 1999; Thomas et al., 2004, Sucháčková et al., 2023). The impacts of global warming on butterfly diversity will likely be mediated by the phylogeographic structure of endangered species and subspecies, including cryptic and symmorphic taxa (Dincă et al. 2011, 2015, Voda et al. 2015). Butterfly genetic lineages are often locally adapted to local climatic regimes, can inhabit particular elevational ranges, or use idiosyncratic habitats and host plant types (Voda et al. 2015, Hill et al. 2021, Polic et al. 2022). Different global warming-induced responses may be expected depending on the phylogeographic distribution of populations, and their degree of genetic isolation, gene flow and introgression patterns between them (Dincă et al. 2013, Talla et al. 2019). Likewise, global-warming induced adaptive trait responses and evolutionary dynamics may be qualitatively different for each genetic lineage (Carnicer et al. 2012).

Despite the importance that underlying phylogeographic structure can have in shaping insect responses to global warming, the studies on the variability of these responses inside taxonomic complexes are still scarce. In this regard, the green-veined white, *Pieris napi* (Linnaeus, 1758), has recently received renewed attention as a model species for the detailed study of global-change induced population declines in Europe, including thermal exposure, adaptive trait responses and microclimatic variability at population level (Espeland et al. 2007, Carnicer et al. 2019, Günter et al. 2019, 2020; Süess et al. 2022, Pruischer et al. 2022, von Schmalensee et al. 2022, 2023, Vives-Ingla et al. 2023). While it is generally recognised that the *Pieris napi* complex includes multiple taxonomic units, this group currently lacks a robust analysis of the genetic variability across its distribution range in the West Palearctic region. The group is also present in the Nearctic, where it also forms a complex of species that has been studied in greater detail (Chew & Watt 2006; Geiger & Shapiro 1992). Finally, the group is also widely distributed in Asia, populations that have been studied with molecular methods and are key to understanding the relationships of the group worldwide (Ge et al. 2023 and references therein). In general, two main clades seem to exist for the *P. napi* complex: one containing most of the Nearctic species and another one mostly Palearctic. Resolving the taxonomy and phylogeography of this group in Europe is key to correctly interpret published and upcoming results on adaptation, ecology and conservation, as well as to forecast differential responses to climate change at high resolution.

As illustrated in Figure 1A, the *P. napi* complex in the Western Palearctic contains at least five taxa (*napi, adalwinda*, *segonzaci*, *bryoniae* and *balcana,* Table S1, Geiger & Scholl, 1985). However, the taxonomic status of these units is controversial and remains poorly understood. No consistent differences in genitalia have been reported among these taxonomic groups in previous studies (Lorković, 1955). At low to medium altitudes, the taxon *napi* (Linnaeus, 1758) is widespread across most of Europe (Figure 1A, yellow area). In contrast, *bryoniae* (Hübner, [1806]) inhabits higher and colder areas of the Alps, the Carpathians and the Tatras, usually at elevations above 1200 m (Porter, 1997, see dark blue area highlighted in Figure 1A), and its distribution apparently extends into Asia (Tshikolovets, 2011). The taxa *napi* and *bryoniae* display hybrid zones in the overlapping areas of their distributional ranges (Petersen, 1963). The relationship between the populations of *napi* and *bryoniae* in Europe has been previously analyzed (Lorković, 1955; Porter, 1997), although not with genomic methods.

**Figure 1.**
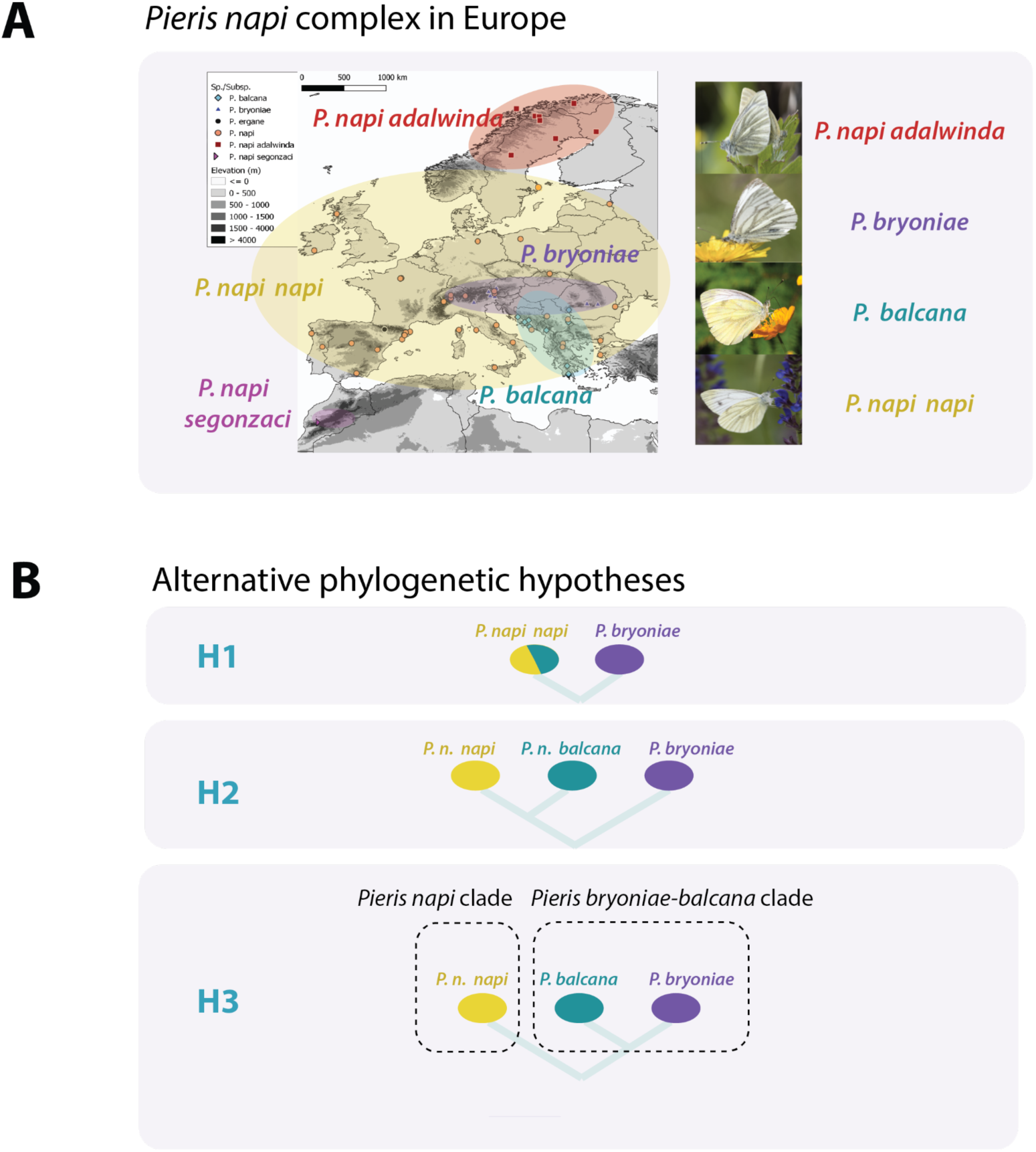
Map showing the distribution of the samples examined in this study, belonging to five taxa in the *Pieris napi* complex: (*napi, adalwinda*, *segonzaci, balcana and bryoniae*).

At much higher latitudes, the taxon *adalwinda* (Fruhstorfer, 1909) inhabits parts of Norway, Sweden and Finland (Petersen, 1947, Figure 1A, red area). Based on morphological differences and the reported contrasting distributional patterns, this taxon is usually considered a subspecies of *P. napi* (Espeland et al., 2007), but other authors have historically hypothesized a putative relationship with *bryoniae* (e.g. Fruhstorfer, 1909). In the northern parts of Scandinavia, *adalwinda* is found all the way from the coastline to the mountains, where typical *napi* is not present (Petersen, 1947). The lowland and montane/arctic forms, both in Scandinavia (*napi* vs. *adalwinda*) and Central Europe (*napi* vs. *bryoniae*), morphologically mainly differ in wing patterns, especially in the amount of melanization and the background color of the wing surface of females (Espeland et al., 2007). Finally, the *P. napi* complex includes North African populations as well (Table S1, S2). Confined to the High Atlas at almost 3000 m, the highly endangered taxon *segonzaci* (le Cerf, 1923) occurs in stream-side alpine areas (Figure 1A, violet color area in Morocco). In addition, a few scattered populations of more typical *napi* have been reported at lower altitudes in the Maghreb, but most of them seem to have become extinct (Tarrier & Delacre 2008).

The taxon *balcana* Lorković, [1969] was described from the Balkans, but its status has been debated for a long time. Compared to *napi, balcana* shows similar wing melanism patterns, but putatively with blurred borders along veins on the hindwing underside and other minor morphological differences (Lorković, 1968). However, Lorković (1968) proposed that the taxon *balcana* was not conspecific with the taxon *napi* because reproductive isolation seems to exist between these taxa. Not only this, but this author suggested that, despite the different morphology, *balcana* may be closer to the taxon *bryoniae* because of karyotypic similarities and also because they seem not to be reproductively isolated in laboratory crosses. Other complementary phylogenetic hypotheses have been also presented linking *napi, segonzaci* and *adalwinda* (Geiger and Scholl 1985, Espeland et al. 2007, Eitschberger & Ströhle 1990).

Overall, inferring the phylogeography of this taxonomic complex is required as a first step for a proper assessment of global change impacts at the European scale. Qualitatively different impacts of global warming could be expected across the taxonomic units due to their contrasting melanic wing patterns, elevation and latitudinal ranges and thermal habitats (Appendix S1). We hypothesize that pronounced shifts or losses in genetic diversity could be concentrated at high-altitude sites, inhabited mainly by melanic forms, and in rear-edge, isolated southern populations (Gómez et al., 2015, Vives-Ingla et al. 2023). In line with these ideas, recent studies have described global-warming induced declines in lowland *P. napi napi* butterfly populations over the last three decades in Spain (Carnicer et al. 2019, Vives-Ingla et al 2023). However, improved phylogenetic and phylogeographic studies are required to delimit proper conservation units at the European scale for these taxa, and the analysis of the genetic structure of populations could facilitate the definition of more robust conservation criteria for the endangered subspecies of this group (Vogler & Desalle, 1994, Dincă et al., 2013). In addition, clarifying the relationships among *P. napi* and its close relatives would shed light on their recent biogeographic history, possibly influenced by multiple factors such as glacial cycles, the evolution of their larval host-plant specificity, and the adaptation to regional thermal regimes (Chew, 1995; Chew & Watt, 2006; Wittstock et al., 2004).

This study aimed at unraveling the conundrum of the *P. napi* complex. The specific objectives were: (**O1**) to assess the phylogenetic relationships of the *P. napi* complex by means of the application of genomic (ddRADseq) techniques; (**O2**) to quantify the patterns of population structure and admixture between the taxa in this complex; (**O3**) to apply species delimitation techniques to propose a solid taxonomic hypothesis; and (**O4**) to assess the relative roles of genetic drift and local adaptation in the complex.

## Methods

### Sample collection

Sample information is summarized in the supplementary tables S2-S4. Table S2 indicates the geographical coordinates of the samples (latitude and longitude), elevation (m), country of origin, and attributed subspecies. After sample quality check analyses, we selected 57 samples for the analyses (Linnaeus, 1758), covering all main taxa considered and spanning their geographic distribution in Europe and North Africa (Figure S1). Additionally, we included 2 samples of *Pieris ergane* (Geyer, 1828) as root (Table S3). Butterflies collected from the field were dried, wings were stored separately as vouchers, and bodies were stored in ethanol 99% at −20°C. Sample collection was performed between 2005 and 2018. Further methodological details on sample collection and processing are specified in the Supplementary methods S1 (DNA extraction, ddRADseq library preparation and sequencing). In addition, Table S3 describes the digestion, ligation and amplification thermal conditions applied in ddRADseq analyses. Table S4 specifies for each sample the raw and filtered number of reads obtained in the ddRADseq, as well as the number of clusters, percentage of missing data and number of final loci.

### ddRADseq data analyses and phylogenetic inference (Objective O1)

Alignment, SNP calling, and initial filtering steps were carried out using ipyrad v.0.7.11 (Eaton and Overcast 2020). The ipyrad analyses were repeated for four subgroups or datasets (A, B, C, D), which are summarized in detail in Table S5. Briefly, group A analyses included all the samples, while groups B-C excludad the outgroup (*P. ergane*) and only the ingroup. As data treatment advanced, the group B was split for further population analyses, henceforth referred as dataset C and dataset D which were also analyzed by ipyrad (Table S5). For assembling ddRADseq data sets we used the *de novo* method whereby sequences are assembled without any reference sources and homology is inferred during alignment clustering by sequence similarity using the program VSEARCH (Rognes, Flouri, Nichols, Quince, & Mahé, 2016). In addition, we also used the reference-based method whereby sequences are mapped to a reference genome using Burrows-Wheeler Alignment tool (Li & Durbin, 2010) based on sequence similarity. We used the yet unpublished genome sequence of a *P. napi* which was generously provided by Christopher Wheat lab (University of Stockholm). The supplementary methods S2 section describes all the ipyrad data processing details.

Ipyrad grouped the reads in 6.76×10⁶ clusters, with values ranging between 0.02×10⁶ and 0.18×10⁶ clusters per sample and a mean of 0.07×10⁶ (Table S6). ipyrad recovered 4544 different loci and the number of loci recovered per sample ranged from 198 to 1714, with a mean of 1207 loci (Table S4). Following initial exploratory analyses and quality checks, the final data sets were assembled using a clustering threshold value (c) of 88%, a maximum number of shared polymorphic sites in a locus (p) of 4 and a filtering threshold (m) that excluded all loci with fewer than 13 individuals (corresponding to 15.3%, 15.6%, 22.8 and 50% of the individuals for the data sets A, B, C and D respectively). Varying the clustering threshold had mostly no impact on the topology inferred.

Maximum likelihood (ML) trees were inferred for the datasets A and B using IQ-TREE 1.6.11 software (Minh, Nguyen, & Von Haeseler, 2013) in an unpartitioned framework, utilizing a single concatenated alignment. A ML tree was also inferred for group A with the reference-based method. For IQ-TREE, the best models of nucleotide substitution were tested with ModelFinder (Wong et al., 2017) and the best-fit model was automatically selected according to the Bayesian Information criterion (BIC). To assess the nodal support, 1000 ultrafast bootstrap (UFBoot) replicates were performed (Hoang et al., 2018) with the -bb command. An SH-aLTR test was also applied with 1000 replicates using the command *– altr* (Guindon et al., 2010). Nodes were considered robust if values of UFBoot ≥ 95 and SH-aLRT ≥ 80. UFBoot has been shown to be largely unbiased compared to standard or alternative bootstrap strategies, and SH-aLRT has been shown to be as conservative as standard bootstrap (Toussaint et al., 2018).

We also inferred the species tree using SNAPP from BEAST v2.7 (Bouckaert et al., 2019). For this, we selected four samples with the highest coverage from each taxon and 5,000 randomly selected SNPs from the neutral and unlinked variant dataset. The SNAPP input file was configured without any constraints on topology or timing. Taxa were grouped into taxon sets based on population designations, and a ClusterTree model with UPGMA clustering was used to initialize the tree. Mutation models incorporated gamma-distributed priors on substitution rates to accommodate rate heterogeneity. Operators for tree scaling, node swapping, and coalescence rate adjustments ensured thorough exploration of parameter space during Markov chain Monte Carlo (MCMC) sampling. No time or topology constraints were applied. SNAPP was run for 10,000,000 Markov chain Monte Carlo (MCMC) generations, and convergence was checked using Tracer v1.7.2 (Rambaut et al., 2018), achieving an effective sample size (ESS) of over 521 for the posterior distribution after discarding 10% of the iterations as burn-in. The resulting trees were merged using TreeAnnotator, and the final species tree was visualized in Figtree (http://tree.bio.ed.ac.uk/software/figtree/). Due to low node posterior support values, we finally visualized the most frequent topologies using DensiTree, to highlight the diversity of topological distributions.

### Population structure analyses and admixture proportion inference (Objective O2)

To detect the presence of cryptic population structure in the association mapping context, identifying the actual subpopulations and probabilistically assign individuals, we used a Bayesian clustering approach by means of the software STRUCTURE v2.3.4 (Falush, Stephens, & Pritchard, 2007). It assumes a model of K populations (i.e., genetic clusters), each of one is characterized by a set of allele frequencies at each locus (Falush et al., 2007).

Using the unlinked SNP frequency data obtained from datasets B, C and D by ipyrad from *de novo* method, the composition of populations was calculated with STRUCTURE (K = number of clusters). The selected burn-in was 100000, followed by 200000 Markov Chain Monte Carlo (MCMC) replicates run to obtain the cluster data. We used Structure Harvester v0.6.94 (Earl & vonHoldt, 2012), which applies the Evanno’s method, an ad hoc statistic ΔK-based on the rate of change in the log probability of data between successive K values (Evanno, Regnaut, & Goudet, 2005), to accurately detect the uppermost hierarchical level of structure. Eventually, the remaining runs of the elected K’s were combined in one per group with CLUMPP (Jakobsson & Rosenberg, 2007). The plots were visualized in Distruct v1.1 (Rosenberg, 2007) and a map was prepared with QGIS v 3.8.1. Finally, a principal component analysis (PCA) was conducted using the Adegenet package (Jombart, 2008) in R to further investigate genetic patterns among populations.

### Taxonomic arrangement and Genotype-Environment Association Analyses (Objectives O3 and O4)

Species delimitation software (SNAPP) was applied to the data for species trees and species delimitation analyses (Leaché and Bouckaert 2018). SNAPP infers species trees under a multispecies coalescent model, estimating species trees directly from biallelic markers (SNP data) without sampling the gene trees at each locus. The method calculates the probability of allele frequency change across ancestor/descendent nodes (Leaché and Bouckaert 2018).

In addition, we employed redundancy analysis (RDA) to investigate multilocus signatures of thermal selection in our focal subspecies complex (Forester et al. 2016; Cheek et al. 2022). RDA analyses model multivariate genetic data as a function of linear combinations of climatic predictor variables by combining multivariate linear regression and PCA (Legendre & Legendre 2012; Cheek et al. 2022). RDA has been proposed as a reliable method to detect subtle, multilocus selection signals of selection due to its low false positive and high true positive rates (Forester et al. 2018; Cheek et al. 2022). However, two RDA methodological challenges may complicate inferences of adaptation across geographic scales. First, genetic structure confounding (Excoffier et al. 2009a,b) can lead to neutral genetic patterns that mirror those of adaptive divergence, particularly when historical isolation and drift have been strong. Second, demographic expansion along latitudinal axes may result in gradual shifts in neutral allele frequencies that parallel environmental gradients, thereby producing misleading signals of selection. To address these confounding factors, we applied both simple and partial RDA (Van Den Wollenberg 1977; Legendre & Legendre 1998), as implemented in the R package vegan (Oksanen et al. 2016).

Partial RDA allows for the explicit partitioning of genetic variance attributable to environmental variables, population structure, and their shared effects, thereby distinguishing true adaptive associations from neutral processes (Capblancq et al. 2021). We first compiled climatic data from the BioClim database (Hijmans et al. 2005), selecting annual mean temperature (BIO1), annual precipitation (BIO12), maximum temperature of the warmest month (BIO5), mean temperature of the warmest quarter (BIO10), precipitation of the warmest quarter (BIO18), isothermality (BIO3), temperature seasonality (BIO4), and precipitation seasonality (BIO15) on the basis of their ecological relevance to this species complex. We additionally derived two vegetation metrics, NDVI and EVI, from MODIS satellite imagery (https://modis.gsfc.nasa.gov), and incorporated high-resolution elevation data from the NASA Shuttle Radar Topographic Mission (https://www.earthdata.nasa.gov) to account for topographic influences. All variables were centered and standardized prior to analysis.

To minimize overfitting in our multivariate model, we implemented a forward selection procedure using forward.sel from the R package packfor (Dray et al. 2009). We adopted a strict α = 0.01 significance threshold for each tested predictor and constrained the global adjusted R² to match that of the full model. Variance inflation factors (VIF) were then evaluated to remove collinear variables. This process yielded two retained predictors: maximum temperature of the warmest month (BIO5, hereafter “temperature”) and precipitation of the warmest quarter (BIO18, hereafter “precipitation”). We conducted a simple RDA with these retained predictors as explanatory variables and the SNP matrix (including candidate adaptive and neutral loci) as response variables. We then performed a partial RDA to control for neutral demographic structure by conditioning on the first two principal components (PC1 and PC2) of a PCA conducted on the same SNP matrix in PLINK2 (Chang et al. 2015). These components were derived after removing candidate SNPs under selection using BayeScan.

Statistical significance for the RDA was evaluated with 1,000 permutations at α = 0.01. To further dissect genomic variation, we used variance partitioning (implemented in vegan) to determine how much of the total genetic variation was uniquely explained by environmental variables, uniquely explained by population structure, or jointly explained by both. All analyses were performed in R version 4.3.2, and full scripts are provided in the Supporting Information.

## Results

### Phylogenetic analyses (O1)

The phylogenetic analyses with IQ-Tree recovered a congruent, robust tree of deep-level divergences in *P. bryoniae*, *P. balcana* and *P. n*. *adalwinda*, supporting the phylogenetic hypothesis H3 (Figure 2; Figure S3-S4). The results thus indicated that *P. bryoniae* and *P. balcana* are phylogenetically related, and recovered them in a clade, separated from *P. napi napi, P. n. adalwinda* and *P. n. segonzaci*. Our study therefore provides a new phylogenetic hypothesis for this common and widespread species group in Europe. It is worth noting the remarkable phylogenetic distance between the three melanic taxa *bryoniae*, *adalwinda* and *segonzaci*.

**Figure 2.**
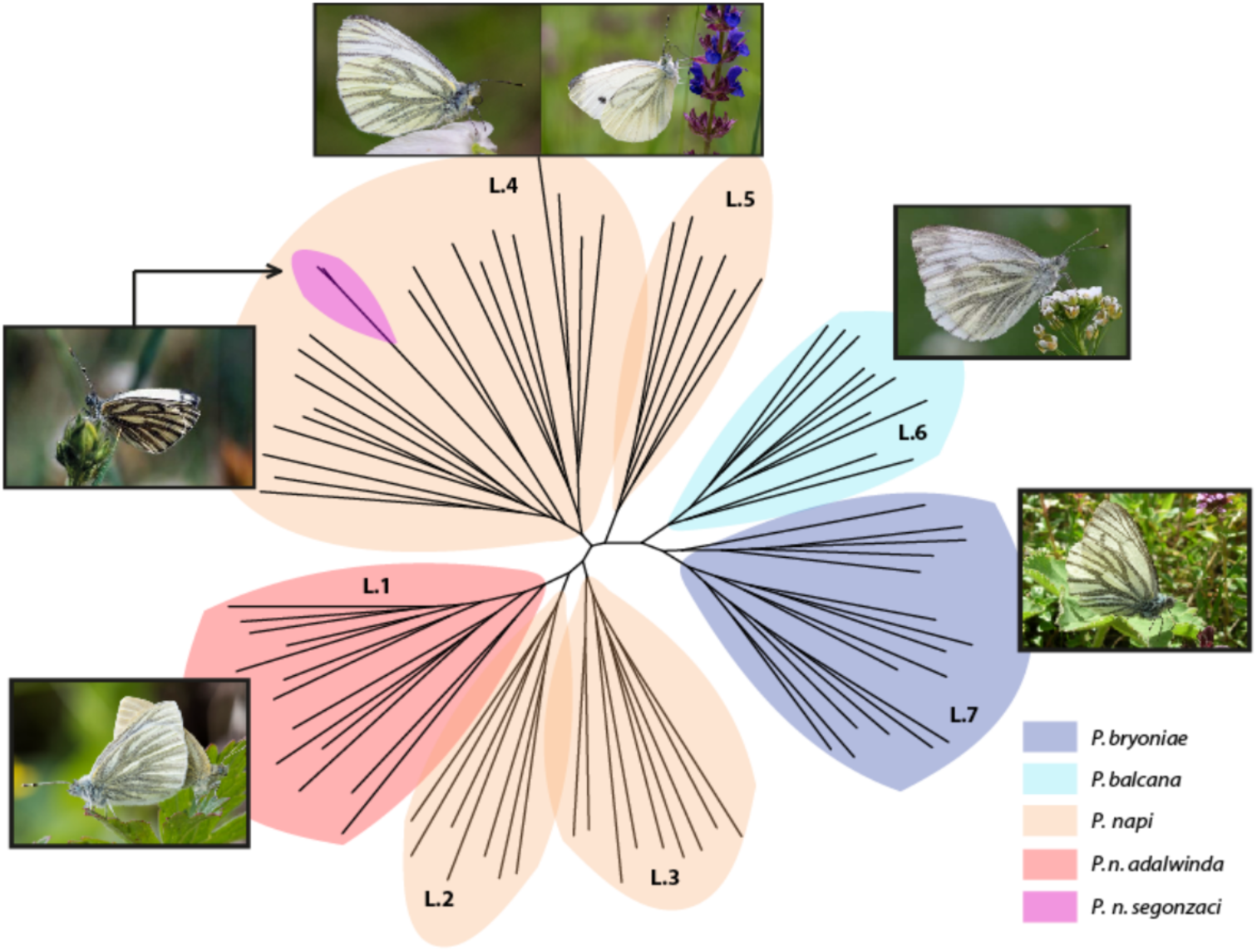
Phylogeny of the *P. napi* complex inferred using ML. The taxa *balcana* and *bryoniae* were recovered as sister clades, which represents a novel taxonomic hypothesis (Figure 1B). The nucleotide substitution model selected in IQ-TREE according to the best information criterion (BIC) was SYM+R3. Support for the phylogenetic nodes and robustness for bootstraps in IQ-TREE analyses is indicated in Figure S3. Nodes were considered robust if support values exceeded 80% in SH-aLRT and 95% in UFBoot. Nodes separating *bryoniae, balcana, adalwinda* and *napi* clades were supported according to these criteria (see Figure S3).

### Population structure and admixture analyses (O2)

PCA analyses (Figure 3) showed two main clusters: PC1 separated specimens within the *napi* clade from species within the *bryoniae-balcana* clade (Figure S6). The two clusters were in line with the previous phylogenetic analyses, and also provided support for the *bryoniae-balcana* clade hypothesis (H3, Figure 1). Each cluster was subdivided in PC2, differentiating *balcana* from *bryoniae* and *adalwinda* from *napi*.

**Figure 3.**
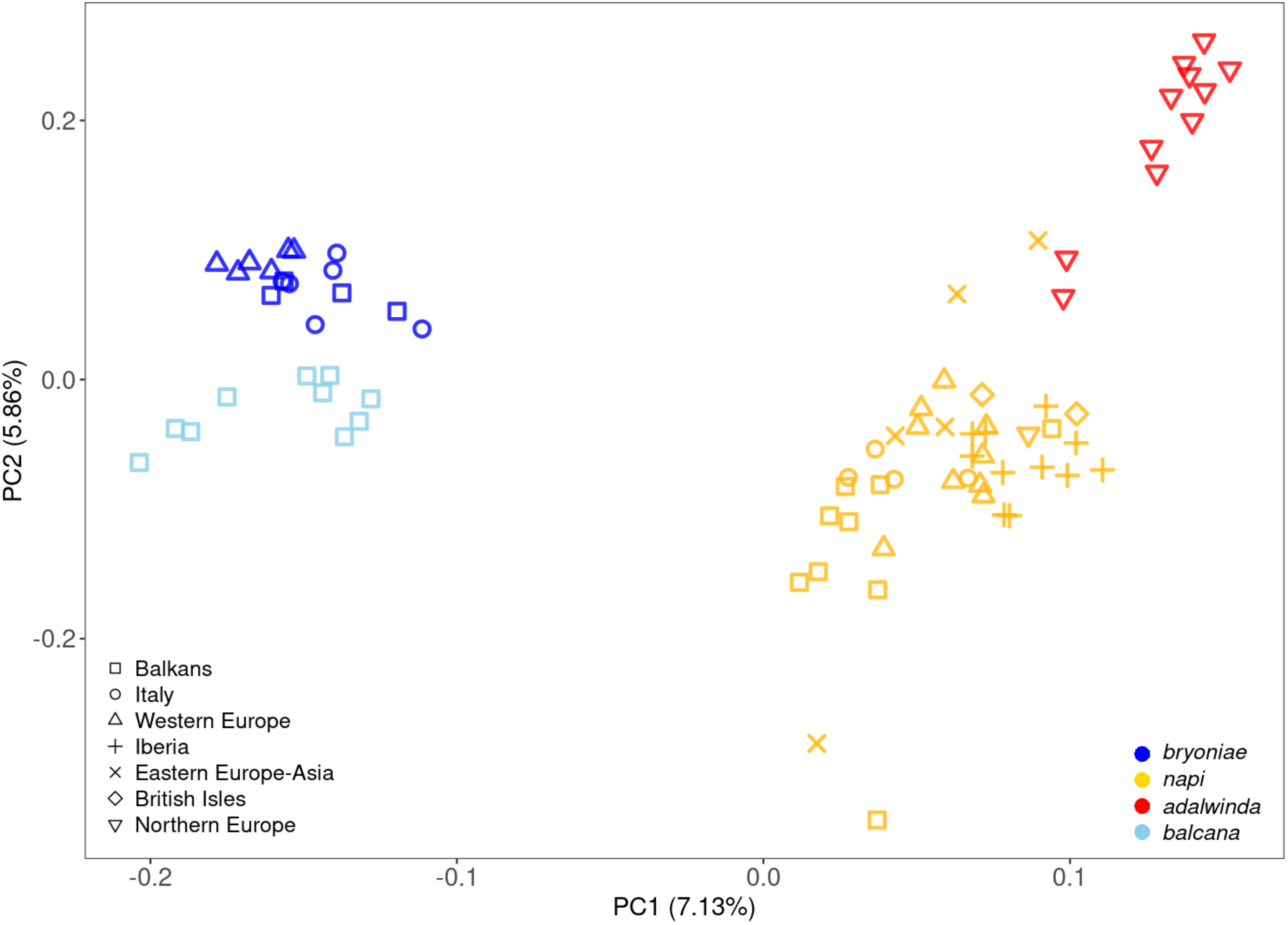
Genetic structure based on a principal components analysis (PCA) of selectively neutral genome-wide SNPs. Marker colours indicate genetic groups corresponding to those shown in the range map (Figure 1A). Shapes represent sampling locations, which are classified into subregions of Europe following the United Nations geoscheme, with Southern Europe further subdivided into the three peninsulas (Iberian, Italian, and Balkan) and the British Isles considered separately. Principal components PC1 and PC2 explain 7.13% and 5.86% of the genetic variation, respectively.

STRUCTURE analyses (Figure 4) yielded an ‘optimal’ clustering K value of 2 according to the ΔK criterion (Figure S4, S5 and Table S76). To better visualize the levels of admixture, we also show the results of K = 5 (ΔK = 0.63). Overall, these analyses reported a hierarchical genetic structure, with a first split between the *napi* and *bryoniae*-*balcana* clades for K = 2 (Figure 3, top), and subsequent division of these two main population groups into other genetic clusters (Figure 3, bottom).

**Figure 4.**
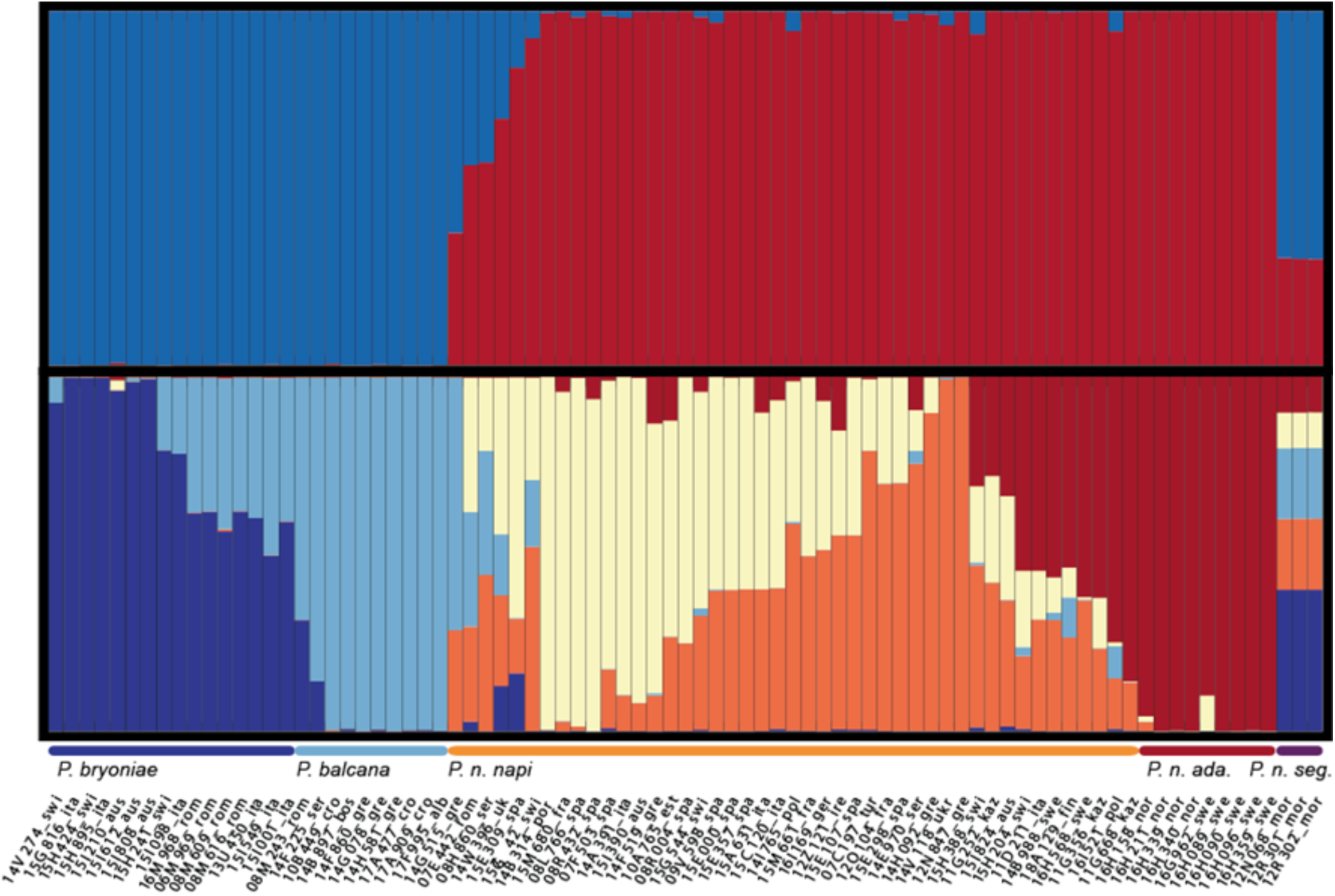
Population structure and admixture inference. Structure analyses are shown for two clustering levels: K=2 (top panel) and K=5 (bottom panel). Each vertical bar represents an individual, with colors corresponding to the proportion of genetic ancestry assigned to each cluster. Geographic variability of the K=5 analysis is detailed in Figure S7. Taxonomic groups are indicated below the bars for reference.

We further analysed each of the two main clades separately with STRUCTURE. For the *P. bryoniae-balcana* clade (i.e. the group of analysis D in table S5), the ‘optimal’ K was 2 (ΔK = 59.42, Table S8). For the *P. napi clade* (group C), the ‘optimal’ K was 6 (ΔK = 5, Table S9). Figure 5 illustrates the geographical distribution of the components and the variation in admixture inference estimated independently for these two major clades.

**Figure 5.**
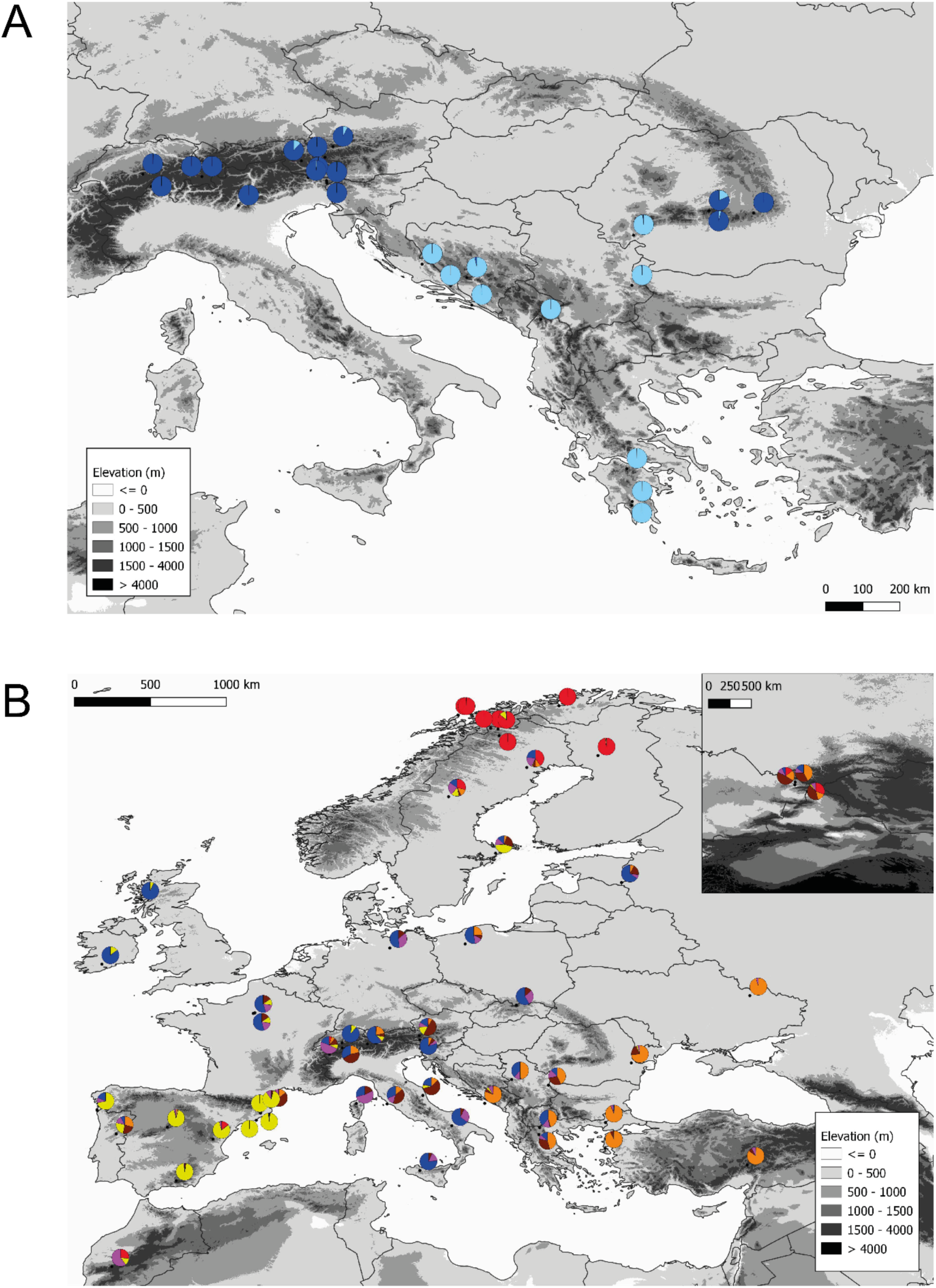
Geographical variation in admixture trends in the two major clades of the *Pieris napi* taxonomic complex in central Europe (Figure 1B). **A**) The *Pieris bryoniae-balcana* clade (K=2; ΔK = 59.42, Table S8). **B**) The *Pieris napi* clade (K=6; ΔK = 5.05, Table S9). Minor dark dots adjacent to the charts represent the exact location of the samples. The upper right corner inset in panel B represents the Kazakhstan samples.

In the *bryoniae-balcana* clade, the considerable proportion of the samples that displayed admixture in Figure 4, depicted by the mixed sky-blue and dark-blue bars, was greatly reduced when this clade was analysed independently (Figure 5A). In this analysis, the two taxa are clearly differentiated and some degree of admixture was only detected in the East of the Alps and in the Carpathians. Based on these results, the distribution of *P. balcana* is shown to encompass all the Balkan peninsula, from Croatia and southern Romania to the Peloponnese. *Pieris bryoniae* is detected in the central and eastern Alps, as well as in the southern Carpathians. In the *napi* clade, most of the samples showed some degree of admixture between lineages, except for the northernmost samples attributable to taxon *adalwinda* (Figure 5B). Different components were most prevalent in different south European peninsula, which suggests a certain degree of differentiation in glacial refugia. These lineages expanded northwards postglacially, mostly from the Italian peninsula. The Kazakhstan samples showed a heterogeneous composition of components, including that of *adalwinda* (inset of Figure 5B). With regards to the taxon *segonzaci*, all the samples showed the same complex clustering composition that is hard to interpret given its isolated distribution.

### Species delimitation analysis (O3)

In line with the previous analyses, SNAPP analyses supported the taxonomic hypothesis H3, i.e. supporting a separate *bryoniae-balcana* clade (Figure 6), and a second clade composed by *P. napi* and *P. n adalwinda*. The main variation between the trees obtained resides in the relationship between *P. napi* and *P. n adalwinda.* Remarkably, this analysis recovers taxon *segonzaci* as the most basal in most of the resulting trees.

**Figure 6.**
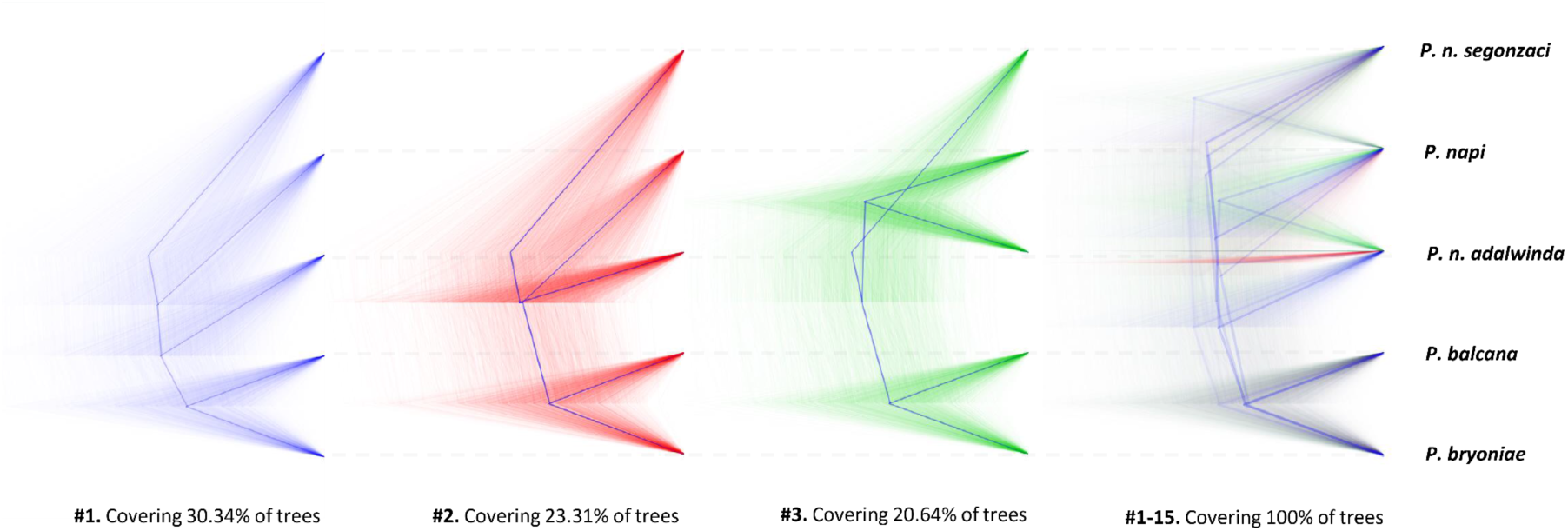
Maximum clade credibility trees generated using SNAPP, summarizing Bayesian posterior probabilities for the phylogenetic relationships among *P. napi* species complex. Each panel represents a distinct topology, with the proportion of posterior trees supporting each topology indicated below (30.34%, 23.31%, and 20.64% for the top three panels, respectively). The final panel (#1–15) includes all observed topologies, summarizing 100% of the posterior distribution. The visualization highlights the variability in phylogenetic relationships across the dataset and emphasizes the dominant topological patterns.

### Analysis of climatic predictor variables in multilocus genetic data (O4)

To assess putative multilocus signatures of natural selection linked to climatic variables, we performed both simple (Figure 7A) and partial (Figure 7B) redundancy analyses (RDA). The simple RDA explained 2.9% (R² = 0.0297) of the total genetic variation (p < 0.001, full model test). Likewise, the partial RDA explained 3.1% (R² = 0.031) of the total genetic variation (p = 0.001, full model test), after accounting for covariates (PC1 and PC1 of a neutral PCA). When we tested for significance by axis using permutation tests (1,000 permutations), the first canonical axis was significant (p < 0.003), while subsequent axes showed progressively lower variance explained and weaker (or non-significant) p-values. We found that temperature and rainfall descriptors explained most of the variation in both types of RDA. The climatic variables highlighted were the maximum temperature of the warmest month and the precipitation of the warmest quarter. Overall, these results indicate that both summer temperature and rainfall descriptors were key factors shaping genetic variation, and that this pattern held true in both the simple and partial RDA frameworks.

**Figure 7.**
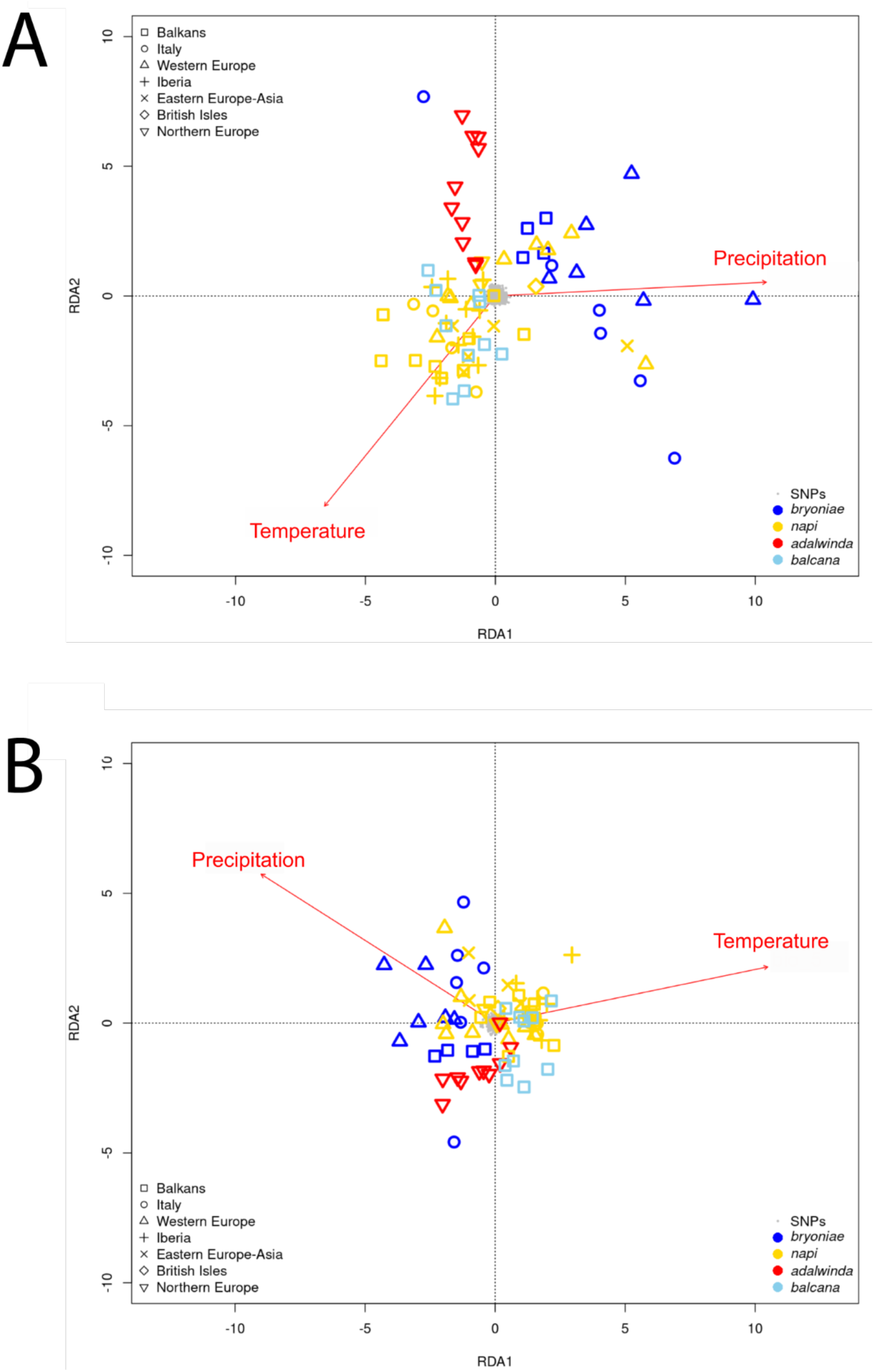
Genetic-environment association analyses in the *Pieris napi* complex. Simple RDA (all genetic variation, **A)** and partial RDA (adaptive genetic variation, **B**) are shown. Colored points represent the projection of individual genotypes on the first two RDA axes. Marker colors correspond to those on the range map in Figure 1. The explanatory variables (maximum temperature of the warmest month and the precipitation of the warmest quarter) are shown within the space defined by RDA1 and RDA2 by labeled vectors. Their contribution to each axis is represented by the length of their orthogonal projections over the scale bars along the top and right sides of the graphs. Arrows indicate the direction of the gradient of variation for the corresponding environmental parameter. The value for each sample point on each explanatory variable can be obtained by an orthogonal projection on the corresponding plotted vector. (**A**) RDA1 and RDA2 axes of a simple RDA based on 88,400 SNPs. (**B**) First two RDA axes of a partial RDA based on 88,400 SNPs conditioned by neutral genetic structure, approximated by the first two principal components of a PCA based on neutral markers. Thus, this plot shows the association between non-neutral genetic signal (presumably under selection) with environmental variables studied.

In the *bryoniae–balcana* clade, the partial RDA (Figure 7B) suggested that *P. balcana* exhibits adaptive variation associated with warmer and drier summer conditions. In contrast, *P. bryoniae* was linked to colder and wetter environments. In the *P. napi* clade, the analyses indicated a predominance of thermal adaptation to colder conditions in *P. n. adalwinda.* Rainfall contributed to some of the observed variability in this clade as well, though to a lesser extent, separating western and eastern populations. The similarity in the trends observed between the simple and partial RDA analyses suggests that climatic factors play an important role in driving adaptive variation in these clades.

## Discussion

### Evolution of the *Pieris napi* complex in Europe: a new phylogeographic hypothesis

We applied ddRADseq techniques to examine phylogeographic patterns in the *Pieris napi* taxonomic complex in Europe. Our analyses unveil the existence of two main evolutionary lineages : the *P. napi* clade and the *P. bryoniae-balcana* clade (Figure 1B, H3). We provide multiple genomic evidence suggesting that *balcana* and *bryoniae* evolved from a common ancestor and are sister taxa, despite their remarkably different morphology and ecology (Figures 2, 3, 5 and 6). This result clarifies the evolutionary origin of the taxon *balcana*, a question that has been debated in previous studies of this taxonomic complex in Europe (Lorković 1962, 1969, Geiger and Shapiro 1992, Porter 1995).

The evolutionary relationships recovered for the rest of the taxa (*napi*, *adalwinda* and *segonzaci*) vary depending on the analyses and are not fully resolved. *Pieris napi* is subdivided into multiple lineages, one of which is the boreal and morphologically differentiated taxon *adalwinda*. All the analyses agree in situating *adalwinda* as a lineage within the *P. napi* clade, but some of the results of SNAPP, suggest a potential relationship to the *P. bryoniae-balcana* clade. This could suggest either some degree of historical admixture between these main clades (not visible in the population structure results) or a contribution by another taxon from Asia. The results for the high-altitude North African taxon *segonzaci* were inconsistent across analyses: from a recent clade within a lineage of *P. napi* in the phylogenetic tree, to a totally admixed entity in STRUCTURE, to a taxon sister to all the rest in SNAPP. These discrepancies are probably due to the low number of specimens and higher missing data for this rare taxon, the status of which should be ideally addressed in a future dedicated study.

Overall, our study indicates that ddRADseq data analyses provide improved descriptions of phylogeographic patterns for closely related cryptic species and also at the subspecies-level, which are possibly relevant for the conservation of endangered cold-adapted forms facing increased global warming pressures, range retraction processes and extreme climatic events in their habitats (IPCC 2022). Our results therefore are in line with other recent genomic studies of multiple insect clades in Europe that have enabled more robust conservation analyses of recently-evolved evolutionary lineages (e.g. Després et al., 2019; Hinojosa et al., 2018; 2019; Hundsdoerfer, Lee, Kitching, & Mutanen, 2019; Jahner et al., 2017; Kozlov, Mutanen, Lee, & Huemer, 2017; Lee et al., 2018; Ryan et al., 2018; Dincă et al., 2019; Lam et al., 2024).

### Varied degrees of admixture between taxa

Previous studies suggested a role for diverging karyotypes and chromosome variability in limiting the gene flow between the clades of the *P. napi* complex. For example, Lorković (1968) reported that in spermatogenesis *P. napi* always showed 25 bivalent chromosomes whereas *bryoniae* and *balcana* were chromosomally more polymorphic. More specifically, he reported for the *bryoniae* and *balcana* clades 25-28 and 26-37 bivalent chromosomes, and between 0 - 5 and 0 - 3 B-chromosomes, respectively. The existence of a hybrid zone of *P. napi* and *P. bryoniae* has been relatively well documented (Lorković, 1955; 1968; Porter, 1997). In this hybrid zone a dominant *napi* phenotype has been reported for hybrid individuals (Lorković, 1955; 1968; Porter, 1997).

In line with these findings, our analyses inferred limited admixture between the *napi* and *bryoniae* in adjacent populations (Figure 4 and S7). Similarly, we found evidence for a reduced level of admixture between *napi* and *balcana*. Considering that a high degree of hybrid sterility between *napi* and *balcana* individuals was reported by Lorković (1968) in crosses, we interpret that this limited degree of admixture could be putatively linked to a consistent and sustained low degree of gene flow and introgression processes between these two clades (Figure 3). In contrast, in the case of the subspecies *adalwinda* (Figure 1) we have inferred a higher degree of admixture with *P. n. napi* populations of the lowland areas of the Scandinavian peninsula, showing a latitudinal clinal trend in admixture (Figure 4, Figure S7). These trends are consistent with previous studies of clinal variability in wing melanism traits and genetic markers in this geographic area (e.g. Espeland 2007, Günter 2019, 2020).

### Phylogeographic refugia and evolutionary divergence

In relation to the southern populations of *P. n. napi*, we identified three major genetic clusters (Figure 3) which geographically overlapped with the three South European Peninsulas (Iberian, Italian and Balkan+Anatolia). This pattern is in agreement with the view of the three major Mediterranean peninsulas of Europe as glacial refugia for multiple taxa (Taberlet et al., 1998; Hewitt 2000, 2001), including butterflies (Schmitt 2007; Dapporto et al., 2024). In line with this idea, our admixture maps (Figures 4, S7) suggest that each of these peninsulas possibly sheltered and produced a different *P. napi* lineage during the glaciations, alike it has occurred in other butterfly complexes such as *Polyommatus coridon/hispana* (Schmitt & Seitz, 2001) and *Melanargia galathea/lachesis* (Habel, Schmitt, & Müller, 2005).

Despite these different glacial refugia and peninsular origins, our results indicate considerable admixture among them as a consequence of postglacial recolonization processes (Figure 4), suggesting that no strong reproductive barriers possibly emerged among the isolated *P. n. napi* refugial lineages in Europe (Geiger and Shapiro 1992). This is consistent with limited reproductive barriers reported in crosses of *P. napi* forms from southern and central European populations in previous studies (Lorković, 1968, Porter et al 1997). The admixture patterns also suggest that, while lineages in all refugia successfully expanded northwards to the mid-latitude areas of Europe, the Italian refugium most notably contributed to the postglacial recolonisation (Figure 4). This result coincides with the average model of the genetic legacy of the Quaternary ice ages for the West Palearctic butterflies (Dapporto et al., 2024). During glacial periods the *adalwinda* lineage most likely shifted south (as the more northern areas north were fully glaciated) and reached central or even southern Europe areas during glacial maxima (Schmitt, 2007). In this case, the admixture was not sufficient as to completely erase the cold-adapted lineage, which would represent an example of a trans-glacial latitudinal layering of populations (Schär et al., 2017). Alternatively, the *adalwinda* lineage could have differentiated in Asia and reached Scandinavia postglacially, as has been proposed for the case of *L. sinapis* (Talla et al., 2019).

Our results also indicate a common ancestor for *P. bryoniae* and *P. balcana*. The geographical and evolutionary origin of this clade remains unclear. Two main geographical hypotheses could be considered: (a) the common ancestor was already a high-mountain specialist, and then the *balcana* lineage adapted to lowland areas, hence, morphological resemblance between *napi* and *balcana* would be due to homoplasy; (b) the common ancestor was a lowland specialist and the *bryoniae* lineage would later speciate occupying alpine regions, therefore, the semblance between *P. napi* and *P. balcana* would be due to plesiomorphy. *Pieris bryoniae* may have speciated as a result of an ecological adaptive process to alpine regions (Lorković, 1955; Porter, 1997). Our results suggest a parallel evolutionary process occurred across lowland mountain range areas in *P. balcana*.

### Species delimitation analyses (SNAPP)

Consistent with admixture and phylogenetic analyses, species delimitation (SNAPP) analyses suggested two major clades in the analyzed European populations, and supported the *bryoniae-balcana* clade (Figure 5). In taxonomy, the discussion about reproductive isolation and the validity of multiple criteria considered to establish the boundaries between species and subspecies has changed over the last decades (de Queiroz, 1998). Presently, different European butterfly species and subspecies may often yield to fertile hybrid descendants. In fact, around 16% of the 496 European butterfly species are known to hybridize. However, the degree of reproductive isolation between the *P. napi* complex subspecies was not experimentally quantified in this study, posing some limitations to the interpretation of the results (Lorković, 1968, Geiger and Shapiro 1992, Porter 1997). In relation to the degree of isolation, the admixture analyses suggested limited gene flow and introgression between the *bryoniae-balcana* clade and the other *P. napi* subspecies (Figure 3). Overall, to properly define the species and subspecies of the *P. napi* complex, further experimental studies on the reproductive isolation of the different subspecies are still possibly warranted, complemented with new phylogeographic studies of the ecological, genetic and evolutionary dynamics of the contact hybrid zones distributed along the European altitudinal and latitudinal gradients (Lorković, 1968, Porter 1997, Espeland et al. 2007, Günter et al 2019, 2020). Previous studies indicate that the *P. balcana* clade presents idiosyncratic karyotypic variability patterns and reproductive isolation in inter-specific crosses (Lorković 1968, Porter 1995, Geiger and Shapiro 1992). In line with these findings, our admixture analyses suggest qualitatively different patterns of gene introgression and reproductive isolation dynamics in relation to both *P. n. napi* populations and *P. bryoniae* populations (Figure 3).

### The *P. napi* complex as a model system to study climate change impacts on insects

We reveal that a similar phenomenon of ecological differentiation linked to climatic adaptation has happened in parallel in the two main clades of the *Pieris napi* complex in Europe: adaptation to summer temperature and precipitation has driven the differentiation of *P. bryoniae* and *P. balcana* (mostly altitudinally), as well as of *P. n. adalwinda* from *P. n. napi* (mostly latitudinally). Such phenomenon is conspicuously mirrored by external morphology: the cold adapted taxa are strongly melanised (presumably improving thermoregulation in cold environments). The case of taxon *segonzaci* in the Maghreb could be added as a third case following the same process.

We suggest that the *Pieris napi* complex in Europe could emerge as a useful model system to study climate-change induced responses in closely related taxa adapted to qualitatively different ecological and latitudinal contexts. The taxon *segonzaci* would provide an opportunity for the study of range retractions and evolutionary processes in rear-edge areas in the Atlas mountains (Northern Africa), a hotspot area for increased climate change impacts (Ali et al. 2022). In central Europe, *P. bryoniae* represents a melanic, cold-adapted species with partial reproductive isolation and dynamic hybrid zones with *P. napi* (Lorković 1968, Porter 1995, Geiger and Shapiro 1992). We here showed that *P. balcana* is a cryptic species, phylogenetically related to *P. bryoniae*, but phenotypically similar to the lowland *P. napi* clades, in terms of wing melanism and climatic preferences. Finally, *P. n. adalwinda* currently constitutes a model subspecies for the study of adaptive responses at high latitudinal, cold, boreal ecosystems, likely presenting low barriers to gene introgression from paler, lowland Scandinavian forms (Espeland et al. 2007, Figure S7).

Over the next decades, range retractions and elevational shifts induced by global warming may likely affect the three high-elevation and melanic forms in this taxonomic complex (i.e. *segonzaci*, *bryoniae* and *adalwinda*). Range retractions could also occur in the paler form *P. balcana,* in populations inhabiting top mountain range areas in the Balkans. In contrast to these taxa, we suggest that the paler *P. n. napi* populations, widely distributed in lowland areas throughout the continent, may possibly expand northwards over the next decades in multiple cold areas of the continent, and increase their altitudinal range limits in multiple European mountain ranges, changing their ecological and evolutionary interactions with the other closely related taxa of the complex. Nevertheless, southern *P. n. napi* populations may also suffer range retraction processes, which could be concentrated in rear edge areas affected by increasing drought and heat wave pressures (Carnicer et al 2019, Vives-Ingla et al. 2023).

### Global warming threats to cold-adapted and cryptic subspecies in the *P. napi* complex: conservation of endangered subspecies

Our analysis suggests that this taxonomic complex possibly contains multiple subspecies that will become progressively endangered by increased global-warming induced pressures. At the European level, 8.5% of the butterfly species (37 species) are considered as threatened, with 0.7% of them being Critically Endangered, 2.8% Endangered and 5% Vulnerable. A further 10% (44 species) of species are classified as Near Threatened (van Swaay et al. 2010, Maes et al. 2019). In the European *P. napi* complex, the Moroccan green-veined white (*Pieris n. segonzaci*), which is restricted to the High Atlas area, stands among the 19 Mediterranean butterfly species highly threatened in the Mediterranean basin (i.e. those butterflies included in the IUCN categories CR, EN or VU). *Pieris n. segonzaci* is one of the four butterfly species categorized as “Vulnerable” (VU) by the IUCN Red list for the Mediterranean basin (Numa et al. 2016). This subspecies is currently affected by habitat degradation due to overgrazing (Numa et al. 2016), and the increase of heat extremes, drought pressures and compound events may currently affect these overgrazed areas (IPCC 2022, Ali et al. 2022). *Pieris n. segonzaci* therefore possibly stands as the most endangered taxon of the analyzed clade.

In contrast, the taxa belonging to the phylogenetically distinct *bryoniae-balcana* clade have been classified as “Least Concern” (LC) in recent IUCN conservation assessments (van Swaay et al. 2010, Maes et al. 2019). Our results could improve the development of conservation assessments, for example allowing the explicit consideration of the unique phylogenetic origin of the *bryoniae-balcana* clade. Range restriction processes and local extinctions could be currently occurring in the populations of *P*. *bryoniae* and *P. balcana,* especially in isolated subpopulations restricted to high-elevation areas. To our knowledge, the possibility of maladaptive gene flow or the introgression of adaptive alleles from other *Pieris napi* subspecies in hybrid zones and across the inhabited elevation gradients and hybrid zones remains poorly assessed (Porter et al. 1997, Valencia-Montoya et al. 2020). Finally, we suggest that *Pieris napi adalwinda* has received a limited degree of attention in recent European conservation assessments (van Swaay et al. 2010, Maes et al. 2019). Our phylogenetic analyses indicate substantial genetic differentiation (Figure 2), suggesting that this subspecies could be ideally included in future conservation assessments of butterflies in Europe.

### Concluding remarks

Our study presents a new phylogenetic hypothesis for the European *Pieris napi* complex. We report a cryptic species case among *P. balcana* and the Balkanic populations of *P. n. napi* and a shift on the *P. napi* complex paradigm, as *P. balcana* emerged as the sister species of *P. bryoniae* (Figure 1b, H3). In addition, we found three main genetic clusters in the European populations of *P. n. napi*, matching with the three major Mediterranean peninsulas of Europe as glacial refugia, plus the notably differentiated boreal taxon *adalwinda*. The present study therefore clarifies the phylogenetic relationships between these taxa and provides a more robust evolutionary background for the detailed study of global-warming induced population declines, adaptive gene introgression dynamics and range retraction processes in cold-adapted forms across Europe (Espeland et al. 2007, Carnicer et al. 2019, Günter et al 2019, 2020; Valencia-Montoya et al. 2020, Süess et al. 2022, Pruischer et al. 2022, von Schmalensee et al. 2022, 2023, Vives-Ingla et al 2023).

