## Supplementary information for "Phylogeography and diversification of the *Pieris napi* species group in the Western Palaearctic"

**S1 Supplementary methods**

**S2 Supplementary methods**

**Supplementary tables S1-S8**

**Supplementary Figures S1-S7**

**S1 Supplementary methods**

*DNA extraction, ddRAD library preparation and Illumina sequencing*

Double-digested RAD-Seq library was prepared following Lee et al. (2018). DNA was extracted from half thoraxes of each specimen using a Qiagen DNeasy Blood & Tissue Kit following the manufacturer’s recommended protocol (Qiagen, Valencia, CA, USA) and stored at -20ºC. DNA was eluted in 100 µl EB buffer and quantified with a multilabel microplate reader (Perkin Elmer Wallac 1420 Victor 2, GMI Inc., Ramsey, Minnesota, USA) using Quant-iT PicoGreen dsDNA assay kit (Thermo Fisher Scientific, Waltham, Massachussets, USA). The mean concentration was 65 ng gDNA / µL with only 11 samples below 20 ng gDNA / µL. Due to low concentrations of gDNA in the original extracts, whole genome amplification (WGA) was performed to all samples using a REPLI-g Mini Kits (Qiagen, Valencia, CA, USA) following the manufacturer’s protocol. Following amplification, a 200 ng of gDNA was digested with PstI and MseI restriction enzymes (New England Biolabs). Following digestion, ligation of double-stranded sequencing adapters was completed in the same tube. The P1 adapter included the Illumina sequencing primer sequences (48 unique 5 bp barcodes), and a TGCA overhang on the top strand to match the sticky end left by PstI. The P2 adapter included the Illumina sequencing primer sequences (same 5 bp barcode), and an AT overhangs on the top strand to match the sticky end left by MseI. It also incorporated a “divergent-Y” to prevent amplification of fragments with MseI cut sites on both ends. Ligated DNA fragments were were purified with 1x AMPure XP magnetic beads (Beckman-Coulter, Brea, CA, USA). Following ligation, size selection was performed by the automated size-selection technology, BluePippin (Sage Science; 2% agarose cartridge). We produced two pooled libraries in four lanes of the machine using automated size selection set to “tight” with a mean of 300 bp. Size selected libraries were eluted in 40 μL volumes and enriched by PCR using library-specific indexed primers complementary to the Illumina paired-end adapters, so that each sample was tagged with a unique combination of barcodes. Amplified DNA fragments were purified again. The quality, size, and concentration of the pooled libraries were finally determined using the MultiNA®(Shimadzu, Kyoto, Japan). The whole process was conducted using Oulu University facilities. Finally, individual fragment libraries were then combined in equimolar amounts and sequenced on an Illumina HiSeq 2500 100 bp paired-end at FIMM (Helsinki, Finland).

**S2 Supplementary methods**

*Ipyrad data processing*

The sequential steps implemented during ipyrad analyses are graphically summarized in Figure S2. During data processing by ipyrad, reads were trimmed of barcodes and adapters and qualitatively filtered using a q-score threshold of 33, with bases below this score converted to Ns (non-informative gap) and any results with more than 5 Ns discarded (Figure S2, step 2). With the collected reads, similar clusters were identified using a threshold of 86% and 88% of similarity (*c*) and were aligned (Figure S2, step 3). Next, we performed joint estimation of heterozygosity and error rate based on a diploid model assuming a maximum of two consensus alleles per individual. We then used the parameters from the previous step, heterozygosity, and error rate, to determine consensus base calls for each allele, and removed consensus sequences with greater than 5 Ns per end of paired-end reads (Figure S2, step 5). Reads of each sample were then clustered and aligned to consensus sequences (Figure S2, step 6). Finally, we filtered the data sets according to a maximum number of indels allowed per read-end of 5, a maximum number of SNPs per locus of 15, a maximum proportion of shared heterozygous sites per locus (*p*) of 2 and 4, and a minimum number of samples per locus (*m*) of 5, 8, 13 and 21 (Figure S2, step 7).

**Supplementary tables S1-S9**

**Table S1.** *D*istributional range and elevation of the taxa treated in this study, belonging to the *Pieris napi* complex.

| **Taxa** | **Elevation** | **Geographical distribution** | **References** |
| --- | --- | --- | --- |
| P. n. napi | In temperate regions below 1200 m. In boreal regions, lower. | All Europe, Mahgreb and unclear ranges in the Middle East and Asia. | Geiger & Scholl, 1985 |
| P. n. adalwinda | From the sea-level, up to the Scandinavian mountains. | North of Scandinavia and at high elevations in Southern Scandinavia | Espeland et al., 2007 |
| P. n. segonzaci | At almost 3000 m. | High Atlas (Morocco) | Higgins, 1999 |
| P. balcana | Samples ranged from 300 m up to almost 1800 m. | Balkans | Lorković, 1968 |
| P. bryoniae | From 1200 up to almost 3000 m. | European Alps, and Carpathians, Tatras and Asia | Porter, 1997 |

**Table S2** Characteristics of the sequenced samples (N= 96 samples).11 samples out of 96 were discarded prior to the obtention of the final alignment and SNP calling as coverage and rate of missing data attained were not suitable for the subsequent analyses. After quality check processes, discarded samples were subsequently not included in the ipyrad and IQ-TREE analyses (groups A, B, C, D, Table S4).

| **Code** | **Presumed sp./subsp.** | **ddRAD sp./subsp.** | **Country** | **Latitude (deg)** | **Longitude (deg)** | **Elevation (m)** | **Quality check** |
| --- | --- | --- | --- | --- | --- | --- | --- |
| **RVcoll11G252** | *P. napi napi* | *P. napi napi* | Kazakhstan | 49.748 | 84.321 | 451 | Approved |
| **RVcoll11G356** | *P. napi napi* | *P. n. adalwinda* | Kazakhstan | 49.379 | 84.263 | 864 | Approved |
| **RVcoll11G668** | *P. napi napi* | *P. n. adalwinda* | Kazakhstan | 48.483 | 85.865 | 1134 | Approved |
| **RVcoll14B988** | *P. napi napi* | *P. napi napi* | Sweden | 59.749 | 18.656 | 41 | Approved |
| **RVcoll16G962** | *P. napi napi* | *P. napi napi* | Sweden | 63.666 | 15.409 | 320 | Approved |
| **RVcoll16H089** | *P. n. adalwinda* | *P. n. adalwinda* | Sweden | 68.407 | 18.318 | 689 | Approved |
| **RVcoll16H090** | *P. n. adalwinda* | *P. n. adalwinda* | Sweden | 68.407 | 18.318 | 689 | Approved |
| **RVcoll16H096** | *P. napi napi* | *P. n. adalwinda* | Sweden | 68.346 | 18.828 | 412 | Approved |
| **RVcoll16H158** | *P. n. adalwinda* | *P. n. adalwinda* | Norway | 69.942 | 23.035 | 61 | Approved |
| **RVcoll16H211** | *P. n. adalwinda* | *P. n. adalwinda* | Norway | 69.942 | 23.035 | 61 | Approved |
| **RVcoll16H339** | *P. n. adalwinda* | *P. n. adalwinda* | Norway | 69.301 | 16.059 | 5 | Approved |
| **RVcoll16H340** | *P. n. adalwinda* | *P. n. adalwinda* | Norway | 69.301 | 16.059 | 5 | Approved |
| **RVcoll16H359** | *P. n. adalwinda* | *P. n. adalwinda* | Sweden | 67.879 | 18.902 | 750 | Approved |
| **RVcoll16H568** | *P. napi napi* | *P. n. adalwinda* | Sweden | 65.679 | 20.813 | 115 | Approved |
| **VDcoll18A129** | *P. n. adalwinda* | *P. n. adalwinda* | Finland | 66.504 | 25.729 | 102 | Approved |
| **RVcoll08M243** | *P. balcana* | *P. balcana* | Romania | 44.866 | 22.417 | 301 | Approved |
| **RVcoll10B449** | *P. balcana* | *P. balcana* | Croatia | 42.820 | 17.680 | 364 | Approved |
| **RVcoll14B897** | *P. balcana* | *P. balcana* | Bosnia & Hrz. | 43.625 | 17.536 | 1229 | Approved |
| **RVcoll14F225** | *P. balcana* | *P. balcana* | Serbia | 43.396 | 22.368 | 583 | Approved |
| **RVcoll14F860** | *P. balcana* | *P. balcana* | Greece | 38.031 | 22.218 | 1786 | Approved |
| **RVcoll14G078** | *P. balcana* | *P. balcana* | Greece | 36.953 | 22.364 | 1655 | Approved |
| **RVcoll14H581** | *P. balcana* | *P. balcana* | Greece | 37.069 | 22.376 | 300 | Approved |
| **RVcoll17A477** | *P. balcana* | *P. balcana* | Croatia | 43.391 | 17.060 | 450 | Approved |
| **RVcoll17A909** | *P. balcana* | *P. balcana* | Croatia | 44.020 | 16.224 | 1174 | Approved |
| **RVcoll17F995** | *P. balcana* | *P. balcana* | Albania | 42.400 | 19.700 | 1400 | Approved |
| **RVcoll14H092** | *P. balcana* | *P. napi napi* | Greece | 40.320 | 21.649 | 833 | Approved |
| **RVcoll14G515** | *P. balcana* | *P. napi napi* | Greece | 40.789 | 21.705 | 930 | Approved |
| **RVcoll14E970** | *P. balcana* | *P. napi napi* | Serbia | 44.114 | 19.765 | 897 | Approved |
| **RVcoll12N847** | *P. balcana* | *P. napi napi* | Greece | 39.359 | 26.234 | 314 | Approved |
| **RVcoll14F519** | *P. balcana* | *P. napi napi* | Greece | 41.140 | 26.244 | 20 | Approved |
| **RVcoll08H860** | *P. balcana* | *P. napi napi* | Serbia | 43.730 | 22.310 | 311 | Approved |
| **RVcoll07E442** | *P. balcana* | *P. napi napi* | Romania | 45.266 | 28.030 | 5 | Approved |
| **RVcoll07C197** | *P. balcana* | *P. napi napi* | Turkey | 38.210 | 36.031 | 1625 | Approved |
| **RVcoll12N068** | *P. n. segonzaci* | *P. n. segonzaci* | Morocco | 31.190 | -7.850 | 2500 | Approved |
| **RVcoll12R301** | *P. n. segonzaci* | *P. n. segonzaci* | Morocco | 31.190 | -7.850 | 2500 | Approved |
| **RVcoll12R302** | *P. n. segonzaci* | *P. n. segonzaci* | Morocco | 31.190 | -7.850 | 2500 | Approved |
| **RVcoll06M968** | *P. bryoniae* | *P. bryoniae* | Romania | 45.517 | 25.929 | 1753 | Approved |
| **RVcoll06M969** | *P. bryoniae* | *P. bryoniae* | Romania | 45.517 | 25.929 | 1753 | Approved |
| **RVcoll08M609** | *P. bryoniae* | *P. bryoniae* | Romania | 45.574 | 24.614 | 1327 | Approved |
| **RVcoll08M616** | *P. bryoniae* | *P. bryoniae* | Romania | 45.585 | 24.629 | 1601 | Approved |
| **RVcoll13U450** | *P. bryoniae* | *P. bryoniae* | Italy | 46.557 | 12.887 | 1870 | Approved |
| **RVcoll14V274** | *P. bryoniae* | *P. bryoniae* | Switzerland | 46.584 | 9.783 | 1905 | Approved |
| **RVcoll15G816** | *P. bryoniae* | *P. bryoniae* | Italy | 46.296 | 8.301 | 1497 | Approved |
| **RVcoll15H241** | *P. bryoniae* | *P. bryoniae* | Switzerland | 46.654 | 8.050 | 1725 | Approved |
| **RVcoll15H424** | *P. bryoniae* | *P. bryoniae* | Switzerland | 46.584 | 9.779 | 1916 | Approved |
| **RVcoll15H895** | *P. bryoniae* | *P. bryoniae* | Italy | 45.740 | 10.853 | 2015 | Approved |
| **RVcoll15I001** | *P. napi napi* | *P. bryoniae* | Italy | 46.415 | 13.440 | 1594 | Approved |
| **RVcoll15I098** | *P. bryoniae* | *P. bryoniae* | Italy | 46.396 | 13.434 | 786 | Approved |
| **RVcoll15I210** | *P. bryoniae* | *P. bryoniae* | Austria | 47.043 | 12.693 | 2420 | Approved |
| **RVcoll15I549** | *P. bryoniae* | *P. bryoniae* | Italy | 46.463 | 13.427 | 1205 | Approved |
| **RVcoll15I612** | *P. bryoniae* | *P. bryoniae* | Austria | 47.460 | 13.618 | 1872 | Approved |
| **RVcoll15I808** | *P. bryoniae* | *P. bryoniae* | Austria | 47.148 | 13.381 | 1746 | Approved |
| **RVcoll09T096** | *P. ergane* | *P. ergane* | Spain | 42.518 | 0.535 | 1541 | Approved |
| **RVcoll14F361** | *P. ergane* | *P. ergane* | Bulgaria | 42.490 | 22.733 | 863 | Approved |
| **RVcoll07F503** | *P. napi napi* | *P. napi napi* | Spain | 41.846 | 2.377 | 726 | Approved |
| **RVcoll08L766** | *P. napi napi* | *P. napi napi* | Spain | 40.821 | -4.009 | 1424 | Approved |
| **RVcoll08R004** | *P. napi napi* | *P. napi napi* | Spain | 40.001 | -0.868 | 1179 | Approved |
| **RVcoll08R432** | *P. napi napi* | *P. napi napi* | Spain | 42.013 | -8.874 | 107 | Approved |
| **RVcoll09V598** | *P. napi napi* | *P. napi napi* | Spain | 37.132 | -3.408 | 1515 | Approved |
| **RVcoll10A765** | *P. napi napi* | *P. napi napi* | Estonia | 57.733 | 27.358 | 119 | Approved |
| **RVcoll11D211** | *P. napi napi* | *P. napi napi* | Italy | 37.838 | 13.429 | 931 | Approved |
| **RVcoll12O104** | *P. napi napi* | *P. napi napi* | France | 42.456 | 9.034 | 631 | Approved |
| **RVcoll12Z121** | *P. napi napi* | *P. napi napi* | Ireland | 52.092 | -8.539 | 227 | Approved |
| **RVcoll14A391** | *P. napi napi* | *P. napi napi* | Italy | 43.023 | 13.627 | 225 | Approved |
| **RVcoll14B314** | *P. napi napi* | *P. napi napi* | Portugal | 40.364 | -7.554 | 1046 | Approved |
| **RVcoll14I765** | *P. napi napi* | *P. napi napi* | Polonia | 49.262 | 20.090 | 1214 | Approved |
| **RVcoll14V118** | *P. napi napi* | *P. napi napi* | Ukraine | 49.917 | 36.217 | 100 | Approved |
| **RVcoll14W396** | *P. napi napi* | *P. napi napi* | Ukraine | 56.540 | -5.800 | 55 | Approved |
| **RVcoll15A631** | *P. napi napi* | *P. napi napi* | Italy | 42.428 | 11.159 | 49 | Approved |
| **RVcoll15C120** | *P. napi napi* | *P. napi napi* | Italy | 40.921 | 15.632 | 779 | Approved |
| **RVcoll15E000** | *P. napi napi* | *P. napi napi* | Spain | 41.302 | 2.124 | 5 | Approved |
| **RVcoll15E107** | *P. napi napi* | *P. napi napi* | Spain | 42.466 | 17.904 | 2150 | Approved |
| **RVcoll15E298** | *P. napi napi* | *P. napi napi* | Spain | 42.221 | 3.094 | 5 | Approved |
| **RVcoll15E309** | *P. napi napi* | *P. napi napi* | Spain | 42.142 | 2.572 | 500 | Approved |
| **RVcoll15E357** | *P. napi napi* | *P. napi napi* | Spain | 41.803 | 2.430 | 1000 | Approved |
| **RVcoll15G544** | *P. napi napi* | *P. napi napi* | Switzerland | 46.298 | 8.064 | 1431 | Approved |
| **RVcoll15G742** | *P. napi napi* | *P. napi napi* | Switzerland | 45.882 | 7.183 | 2202 | Approved |
| **RVcoll15H204** | *P. napi napi* | *P. napi napi* | Switzerland | 46.654 | 8.050 | 1725 | Approved |
| **RVcoll15H388** | *P. napi napi* | *P. napi napi* | Switzerland | 46.576 | 9.803 | 2059 | Approved |
| **RVcoll15I390** | *P. napi napi* | *P. napi napi* | Austria | 46.869 | 13.426 | 1397 | Approved |
| **RVcoll15I824** | *P. napi napi* | *P. napi napi* | Austria | 47.148 | 13.381 | 1746 | Approved |
| **RVcoll15M661** | *P. napi napi* | *P. napi napi* | France | 48.727 | 2.023 | 160 | Approved |
| **RVcoll15M680** | *P. napi napi* | *P. napi napi* | France | 48.688 | 1.919 | 130 | Approved |
| **RVcoll16I169** | *P. napi napi* | *P. napi napi* | Germany | 53.206 | 11.346 | 33 | Approved |
| **RVcoll16I521** | *P. napi napi* | *P. napi napi* | Polonia | 53.487 | 16.553 | 140 | Approved |
| **RVcoll10B334** | *P. balcana* | *-* | Bulgaria | 41.623 | 25.727 | 155 | Discarded |
| **RVcoll12N067** | *P. n. segonzaci* | *-* | Morocco | 31.190 | -7.850 | 2500 | Discarded |
| **RVcoll14E050** | *P. bryoniae* | *-* | Italy | 44.745 | 7.110 | 1976 | Discarded |
| **RVcoll14N469** | *P. bryoniae* | *-* | Ukraine | 48.139 | 24.447 | 1457 | Discarded |
| **RVcoll08L792** | *P. napi napi* | *-* | Spain | 43.019 | -4.344 | 1392 | Discarded |
| **RVcoll14E053** | *P. napi napi* | *-* | Italy | 44.203 | 7.248 | 1531 | Discarded |
| **RVcoll14W968** | *P. napi napi* | *-* | United Kingdom | 50.210 | -3.709 | 69 | Discarded |
| **RVcoll15G161** | *P. napi napi* | *-* | France | 46.365 | 5.898 | 1064 | Discarded |
| **RVcoll15O591** | *P. napi napi* | *-* | France | 42.997 | -0.848 | 616 | Discarded |
| **RVcoll15Q121** | *P. napi napi* | *-* | Russia | 55.661 | 38.263 | 120 | Discarded |
| **RVcoll16H347** | *P. n. adalwinda* | *-* | Norway | 69.300 | 16.059 | 5 | Discarded |

**Table S3.** A summary of the ddRADseq protocol conditions

| Digestion | | |
| --- | --- | --- |
| Step | **Temperature (ºC)** | **Time (min)** |
| Digestion | 37 | 180 |
| Deactivation | 65 | 20 |
| Ligation | | |
| Step | **Temperature (ºC)** | **Time (min)** |
| Ligation | 16 | 180 |
| Amplification I | | |
| Step | **Temperature (ºC)** | **Time (s)** |
| Denaturation I | 98 | 30 |
| Denaturation II | 98 | 20 |
| Annealing | 60 | 30 |
| Elongation I | 72 | 40 |
| Elongation II | 72 | 600 |
| Amplification II | | |
| Step | **Temperature (ºC)** | **Time (min)** |
| Denaturation | 98 | 3 |
| Annealing | 30 | 2 |
| Elongation | 72 | 12 |

**Table S4** Information about the ddRADseq data obtained per sample. The number of reads represents those kept by ipyrad and the number of clusters are those grouped by the same program. Percentage of missing data and final number of loci are from the final filtered loci data set.

| **Code** | **Nº reads (raw)** | **Nº read (filtered)** | **Nº clusters** | **Missing data (%)** | **Nº final loci** |
| --- | --- | --- | --- | --- | --- |
| **RVcoll06M968** | 1501151 | 1499915 | 26876 | 77.3 | 1031 |
| **RVcoll06M969** | 3343708 | 3340866 | 49503 | 78.0 | 1000 |
| **RVcoll08M609** | 5312147 | 5307584 | 89327 | 70.4 | 1346 |
| **RVcoll08M616** | 6873887 | 6867899 | 118215 | 65.7 | 1559 |
| **RVcoll13U450** | 5565696 | 5560983 | 116412 | 71.8 | 1282 |
| **RVcoll14V274** | 6935084 | 6928955 | 97281 | 75.2 | 1125 |
| **RVcoll15G816** | 1752075 | 1750702 | 103812 | 69.5 | 1385 |
| **RVcoll15H241** | 2851582 | 2849213 | 69532 | 71.6 | 1290 |
| **RVcoll15H424** | 719794 | 719183 | 60842 | 74.4 | 1163 |
| **RVcoll15H895** | 1388798 | 1387613 | 58248 | 70.1 | 1360 |
| **RVcoll15I098** | 2496822 | 2494703 | 70899 | 70.1 | 1359 |
| **RVcoll15I210** | 4515317 | 4511315 | 116669 | 67.5 | 1477 |
| **RVcoll15I549** | 1605713 | 1604370 | 79634 | 69.5 | 1386 |
| **RVcoll15I612** | 2076133 | 2074303 | 58367 | 71.2 | 1308 |
| **RVcoll15I808** | 7311344 | 7305166 | 76976 | 74.7 | 1148 |
| **RVcoll09T096** | 1802852 | 1801417 | 98690 | 93.5 | 295 |
| **RVcoll14F361** | 1301475 | 1300337 | 25454 | 95.6 | 198 |
| **RVcoll15A631** | 1342864 | 1341869 | 40170 | 82.7 | 787 |
| **RVcoll15C120** | 1596657 | 1595314 | 34629 | 79.5 | 932 |
| **RVcoll07F503** | 1154409 | 1153431 | 68927 | 76.0 | 1091 |
| **RVcoll10A765** | 2058470 | 2056703 | 99277 | 69.0 | 1409 |
| **RVcoll11D211** | 2284342 | 2282482 | 61434 | 76.5 | 1070 |
| **RVcoll11G252** | 273003 | 272783 | 21157 | 83.8 | 738 |
| **RVcoll11G356** | 1005271 | 1004433 | 32859 | 76.7 | 1060 |
| **RVcoll11G668** | 781050 | 780392 | 54281 | 74.5 | 1157 |
| **RVcoll12O104** | 5525689 | 5521331 | 108386 | 72.2 | 1263 |
| **RVcoll12Z121** | 607152 | 606663 | 29816 | 79.3 | 939 |
| **RVcoll14A391** | 978849 | 978063 | 38106 | 77.9 | 1003 |
| **RVcoll14B314** | 4283998 | 4280449 | 90559 | 72.6 | 1243 |
| **RVcoll14I765** | 3930323 | 3927211 | 65934 | 72.8 | 1237 |
| **RVcoll15I001** | 14346343 | 14334281 | 90701 | 82.9 | 775 |
| **RVcoll15I390** | 1739580 | 1738097 | 63809 | 74.6 | 1156 |
| **RVcoll15I824** | 859667 | 858958 | 41275 | 85.8 | 646 |
| **RVcoll15M661** | 3450957 | 3448044 | 90635 | 71.7 | 1287 |
| **RVcoll15M680** | 7520537 | 7513447 | 95027 | 72.2 | 1264 |
| **RVcoll16H158** | 4246566 | 4242939 | 123959 | 67.5 | 1475 |
| **RVcoll16H211** | 3453190 | 3450150 | 88954 | 67.1 | 1495 |
| **RVcoll16H339** | 1295393 | 1294285 | 36829 | 73.3 | 1214 |
| **RVcoll16H340** | 1769617 | 1768313 | 96961 | 70.1 | 1357 |
| **RVcoll16I169** | 1067877 | 1067015 | 49340 | 73.5 | 1202 |
| **RVcoll16I521** | 5695727 | 5691646 | 62208 | 73.4 | 1208 |
| **RVcoll08L766** | 6377550 | 6371940 | 129524 | 65.9 | 1550 |
| **RVcoll08R004** | 1479256 | 1477929 | 53819 | 74.7 | 1151 |
| **RVcoll08R432** | 1592583 | 1591247 | 45879 | 75.8 | 1100 |
| **RVcoll09V598** | 8864257 | 8856917 | 181043 | 62.3 | 1714 |
| **RVcoll14B988** | 2318792 | 2316911 | 51198 | 79.0 | 954 |
| **RVcoll14V118** | 1255286 | 1254236 | 44255 | 80.8 | 873 |
| **RVcoll14W396** | 2953849 | 2951234 | 97861 | 70.4 | 1344 |
| **RVcoll15E000** | 3853046 | 3849943 | 104230 | 66.9 | 1503 |
| **RVcoll15E107** | 3214579 | 3211904 | 127729 | 66.3 | 1532 |
| **RVcoll15E298** | 1535831 | 1534591 | 62992 | 74.3 | 1168 |
| **RVcoll15E309** | 1995029 | 1993234 | 65303 | 67.9 | 1460 |
| **RVcoll15E357** | 2405451 | 2403268 | 79937 | 68.0 | 1453 |
| **RVcoll15G544** | 2051726 | 2050018 | 95020 | 73.2 | 1218 |
| **RVcoll15G742** | 3730423 | 3727290 | 59312 | 72.8 | 1234 |
| **RVcoll15H204** | 887311 | 886618 | 38996 | 74.6 | 1154 |
| **RVcoll15H388** | 2496492 | 2494404 | 78885 | 70.4 | 1345 |
| **RVcoll16G962** | 1655340 | 1653823 | 80115 | 69.2 | 1400 |
| **RVcoll16H089** | 2518625 | 2516347 | 106867 | 70.2 | 1354 |
| **RVcoll16H090** | 1805277 | 1803773 | 70150 | 73.3 | 1212 |
| **RVcoll16H096** | 863546 | 862817 | 58692 | 73.1 | 1223 |
| **RVcoll16H359** | 6117915 | 6112379 | 127306 | 69.6 | 1380 |
| **RVcoll16H568** | 7398916 | 7392605 | 136147 | 71.5 | 1295 |
| **VDcoll18A12** | 2384865 | 2382798 | 88999 | 70.8 | 1329 |
| **RVcoll07C197** | 1170115 | 1169133 | 59635 | 74.9 | 1139 |
| **RVcoll07E442** | 4231369 | 4227664 | 152356 | 63.3 | 1669 |
| **RVcoll08H860** | 1443568 | 1442388 | 86055 | 72.1 | 1266 |
| **RVcoll08M243** | 1332237 | 1331074 | 70553 | 68.9 | 1414 |
| **RVcoll10B449** | 731386 | 730723 | 57246 | 69.1 | 1406 |
| **RVcoll12N847** | 1575709 | 1574375 | 60492 | 73.2 | 1220 |
| **RVcoll14B897** | 3912913 | 3909632 | 88332 | 68.6 | 1428 |
| **RVcoll14E970** | 1581765 | 1580409 | 63744 | 75.8 | 1098 |
| **RVcoll14F225** | 2335177 | 2333276 | 81045 | 71.2 | 1307 |
| **RVcoll14F519** | 1623473 | 1622049 | 61774 | 73.9 | 1186 |
| **RVcoll14F860** | 3547528 | 3544695 | 131588 | 66.1 | 1541 |
| **RVcoll14G078** | 5098832 | 5094790 | 168112 | 69.2 | 1400 |
| **RVcoll14G515** | 2265017 | 2263254 | 74639 | 72.0 | 1274 |
| **RVcoll14H092** | 5726441 | 5721563 | 88775 | 73.1 | 1224 |
| **RVcoll14H581** | 4714884 | 4711122 | 163422 | 66.4 | 1527 |
| **RVcoll17A477** | 1594506 | 1593145 | 57990 | 67.3 | 1486 |
| **RVcoll17A909** | 1850454 | 1848882 | 82644 | 67.5 | 1478 |
| **RVcoll17F995** | 7517767 | 7510938 | 177632 | 66.3 | 1533 |
| **RVcoll12N068** | 1086875 | 1085993 | 27081 | 91.6 | 381 |
| **RVcoll12R301** | 1245664 | 1244586 | 39359 | 87.6 | 562 |
| **RVcoll12R302** | 15274416 | 15260873 | 90764 | 90.9 | 414 |

**Table S5.** Summary of the four data sets used for the *ddRADseq* data analyses. Dataset B was split in two main groups (datasets C and D) suggested by STRUCTURE and phylogenetic analysis (corresponding respectively to taxa within *P. napi clade* and *P. bryoniae-balcana clade*).

| **Dataset** | **Number of samples** | **Outgroup** | **P. napi** | **P. bryoniae** |
| --- | --- | --- | --- | --- |
| **A** | 85 | YES | YES | YES |
| **B** | 83 | NO | YES | YES |
| **C** | 57 | NO | YES | NO |
| **D** | 26 | NO | NO | YES |

**Table S6.** Summary statistics of the datasets filtered and aligned by ipyrad.

| **Data set** | **Length of the concatenation (bp)** | **Number of unlinked SNP** | **Number of loci** | **Percentage of missing data** | **Number**  **of variable sites** | **Number of parsimony informative sites** |
| --- | --- | --- | --- | --- | --- | --- |
| **A** | 793017 | 4465 | 4544 | 73.43% | 48232 (6%) | 23923 (3%) |
| **B** | 801806 | 4512 | 4595 | 72.69% | 48363 (6%) | 24146 (3%) |
| **D** | 546968 | 3088 | 3129 | 62.21% | 33778 (6%) | 16783 (3%) |
| **C** | 356132 | 1966 | 2024 | 27.73% | 18752 (5%) | 9279 (2%) |

**Table S7.** Evanno table provided by Structure Harvester using the unlinked SNP’s matrix produced by the *de novo* method and dataset B.

| **# K** | Reps | Mean LnP(K) | Stdev LnP(K) | Ln'(K) | \|Ln''(K)\| | **Delta K** |
| --- | --- | --- | --- | --- | --- | --- |
| **1** | 10 | -61714.65 | 12.70 | NA | NA | **NA** |
| **2** | 10 | -58047.88 | 25.00 | 3666.78 | 1611.21 | **64.45** |
| **3** | 10 | -55992.31 | 194.46 | 2055.57 | 481.49 | **2.48** |
| **4** | 10 | -54418.22 | 237.40 | 1574.09 | 52.28 | **0.22** |
| **5** | 10 | -52896.41 | 579.95 | 1521.81 | 363.19 | **0.63** |
| **6** | 10 | -51737.79 | 564.21 | 1158.62 | 216.56 | **0.38** |
| **7** | 10 | -50795.73 | 1308.17 | 942.06 | 332.03 | **0.25** |
| **8** | 10 | -49521.64 | 1227.95 | 1274.09 | 309.08 | **0.25** |
| **9** | 10 | -48556.63 | 1172.46 | 965.01 | 265.29 | **0.23** |
| **10** | 10 | -47856.91 | 2596.13 | 699.72 | 11149.35 | **4.29** |
| **11** | 10 | -58306.54 | 44501.40 | -10449.63 | 22483.13 | **0.51** |
| **12** | 10 | -46273.04 | 2065.36 | 12033.50 | 25977.01 | **12.58** |
| **13** | 10 | -60216.54 | 35366.24 | -13943.51 | 6130.29 | **0.17** |
| **14** | 10 | -80290.34 | 63791.93 | -20073.79 | 36639.57 | **0.57** |

**Table S8.** Evanno table provided by Structure Harvester using the unlinked SNP’s matrix produced by the *de novo* method for the *Pieris bryoniae-balcana* clade (group D).

| **# K** | **Reps** | **Mean LnP(K)** | **Stdev LnP(K)** | **Ln'(K)** | **\|Ln''(K)\|** | **Delta K** |
| --- | --- | --- | --- | --- | --- | --- |
| 1 | 10 | -22391.64 | 10.23 | NA | NA | **NA** |
| 2 | 10 | -21460.12 | 8.94 | 931.52 | 531.17 | **59.42** |
| 3 | 10 | -21059.77 | 31.26 | 400.35 | 149.51 | **4.78** |
| 4 | 10 | -20808.93 | 320.33 | 250.84 | 99.66 | **0.31** |
| 5 | 10 | -20458.43 | 50.90 | 350.50 | 210.72 | **4.14** |
| 6 | 10 | -20318.65 | 555.65 | 139.78 | 112.67 | **0.20** |
| 7 | 10 | -20066.19 | 338.85 | 252.45 | NA | **NA** |

**Table S9.** Evanno table provided by Structure Harvester using the unlinked SNP’s matrix produced by the *de novo* method for the *Pieris napi* clade (group C, table S5)

| **# K** | Reps | Mean LnP(K) | Stdev LnP(K) | Ln'(K) | \|Ln''(K)\| | **Delta K** |
| --- | --- | --- | --- | --- | --- | --- |
| **1** | 10 | -39420.77 | 17.01 | NA | NA | **NA** |
| **2** | 10 | -37663.01 | 230.29 | 1757.77 | 1027.69 | **4.46** |
| **3** | 10 | -36932.93 | 1874.36 | 730.08 | 795.53 | **0.42** |
| **4** | 10 | -35407.32 | 571.92 | 1525.61 | 1313.38 | **2.30** |
| **5** | 10 | -35195.09 | 2813.34 | 212.23 | 1430.74 | **0.51** |
| **6** | 10 | -33552.13 | 259.75 | 1642.97 | 1312.91 | **5.05** |
| **7** | 10 | -33222.07 | 1231.19 | 330.05 | 901.40 | **0.73** |
| **8** | 10 | -31990.62 | 254.13 | 1231.45 | 947.31 | **3.73** |
| **9** | 10 | -31706.48 | 988.73 | 284.14 | NA | **NA** |

**Supplementary figures S1-S7**

**Figure S1.** Geographical distribution of the *Pieris* samples used in the analyses.

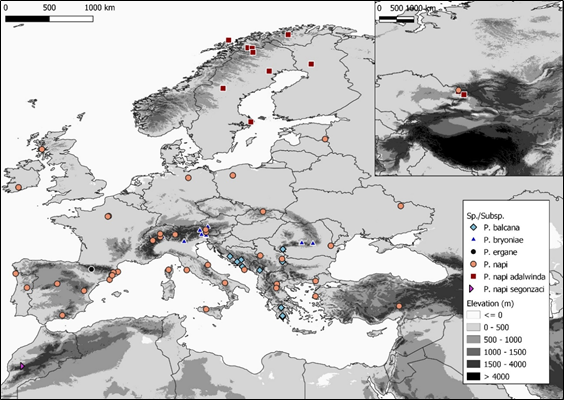

**Figure S2.** A diagram summarizing all the sequential steps implemented in the ipyrad analyses. Source: https://ipyrad.readthedocs.io/

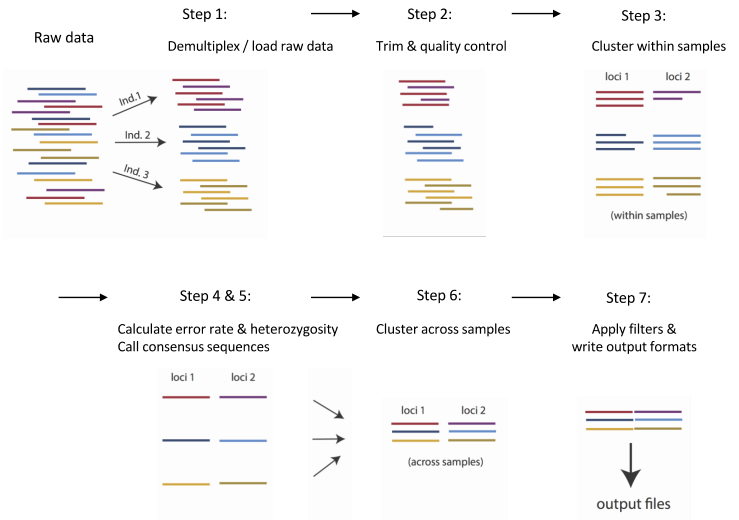

**Figure S3** Phylogeny of the *Pieris napi complex* (ML) with outgroup species (*P. ergane*). The patterns were consistent with the analyses reported in Figure 2 (ML phylogeny without outgroup). The results support the taxonomic hypothesis H3 (Figure 1B). The nucleotide substitution model selected in IQ-TREE according to BIC was TVMe+R3. Nodes are considered robust if support values exceed 80% in SH-aLRT and 95% in UFBoot. Robustness for both bootstraps is indicated with ***,** +/- for SH-aLRT, and -/+ for UFBoot. The tree phylogeny was inferred based on IQ-TREE align (*de novo* method). Images: Vlad E. Dincă.
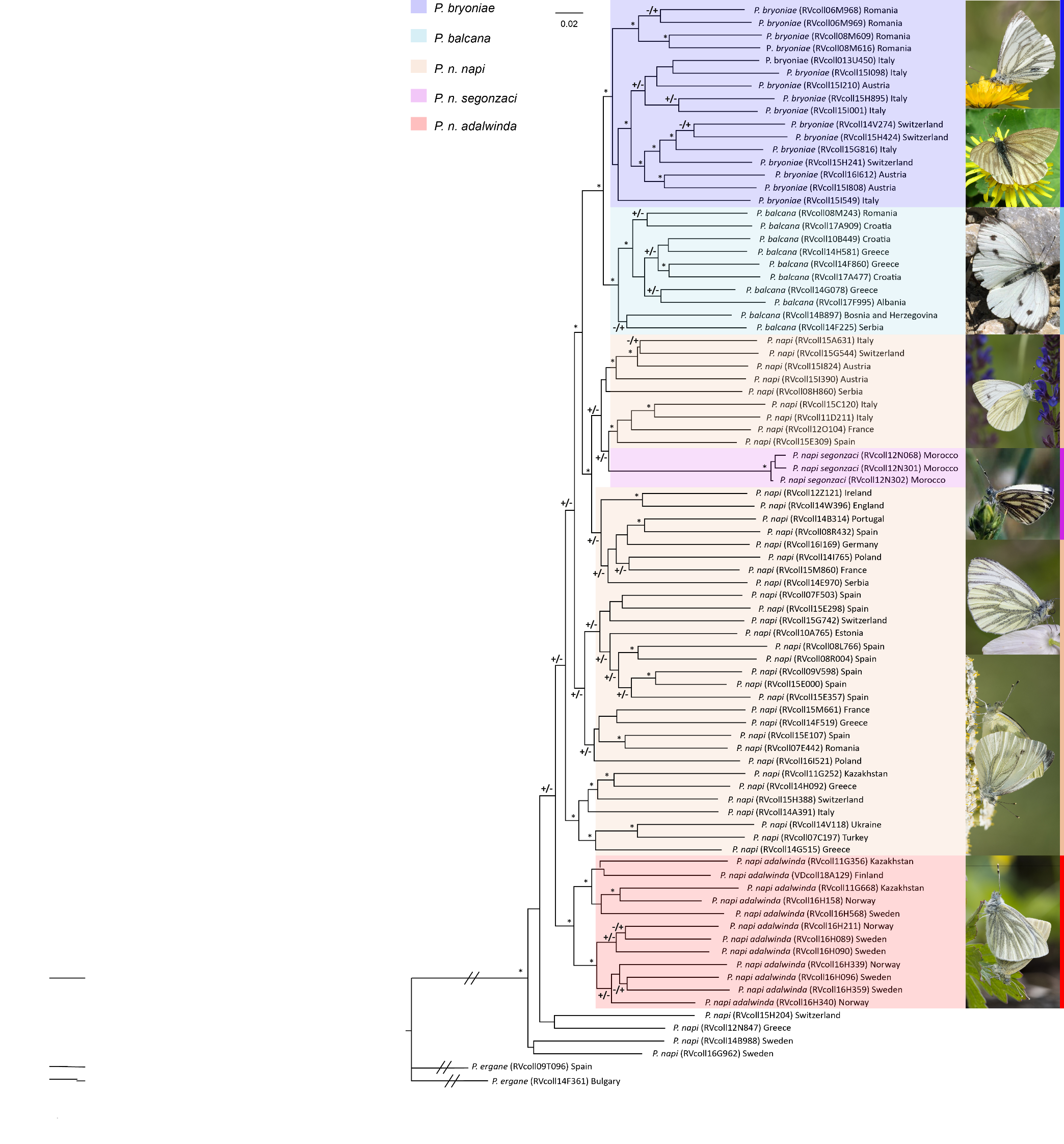

**Figure S4.** Variation of ΔK according to the number of clusters (#K) from STRUCTURE analyses. The *Delta K* statistic identifies the K with the highest likelihood and lowest variance.

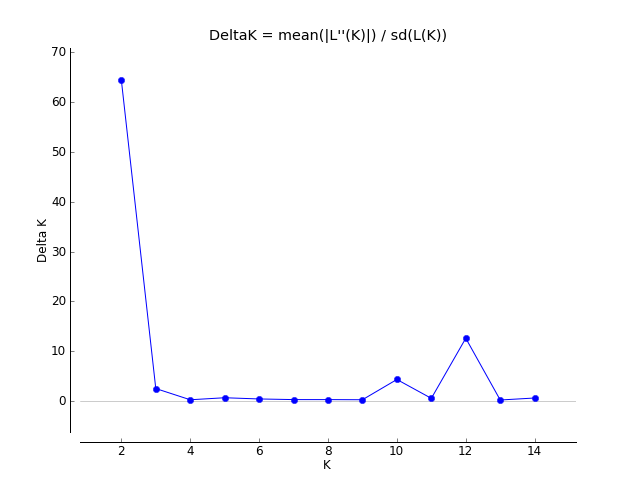

**Figure S5.** Population admixture analyses based on ddRADseq *de novo* assemblies. STRUCTURE analyses for K = 3 (∆K = 2.3).

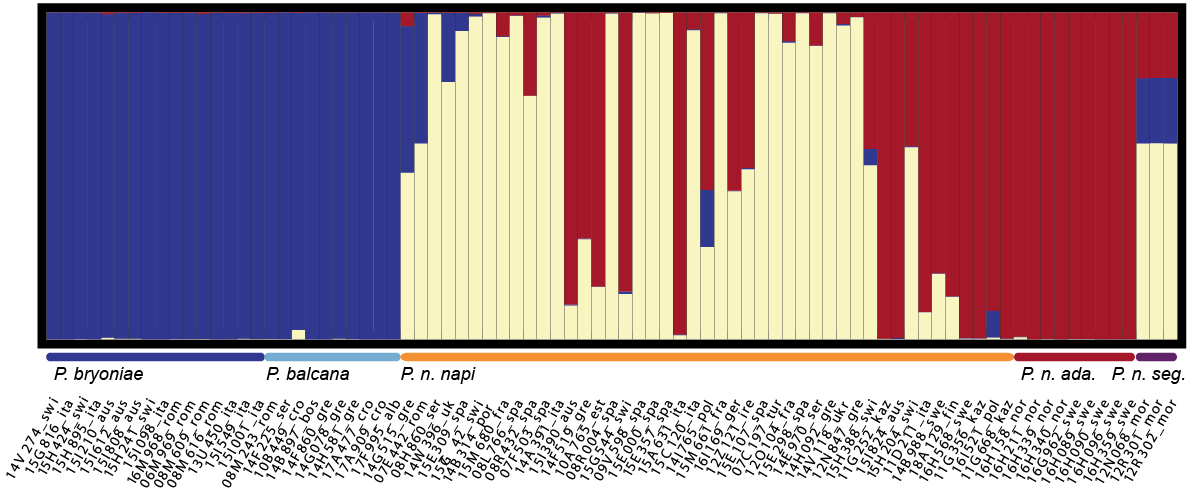

**Figure S6.** PCA analysis of the genetic variation of ddRADSeq de novo alignment (Group B).

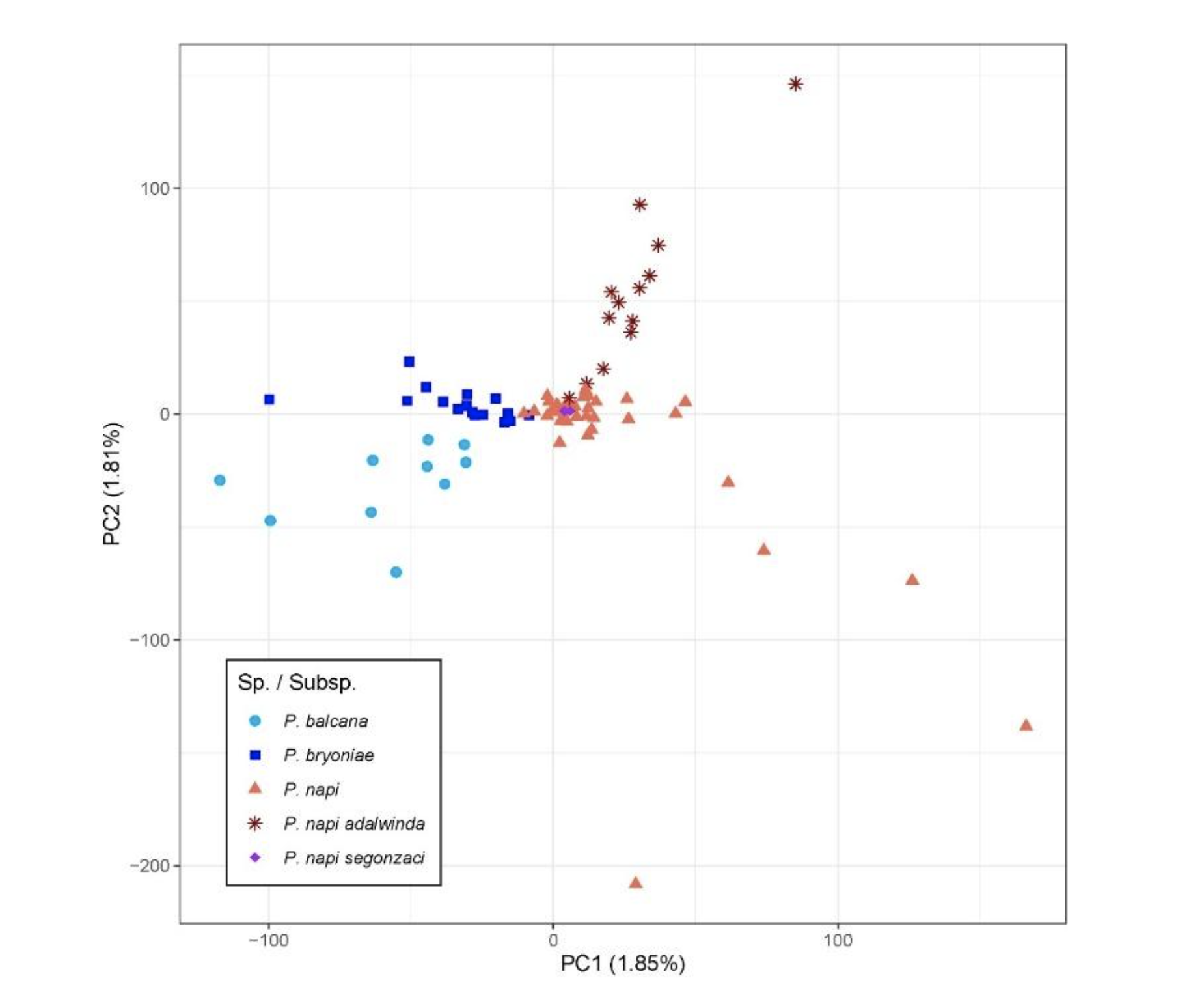

**Figure S7.** European map showing the pie charts of each sample’s probable assignment to five different populations in admixture analyses (K=5 in STRUCTURE) for all the *Pieris napi* complex samples analyzed (Group B, Table S4). Minor dark dots adjacent to the charts represent the exact location of the samples. The upper right corner inset represents the Kazakhstan samples. *P. napi* were approximately grouped in two longitudinal clusters, showing a yellow (West) to orange (East) shifting pattern. *P. n. adalwinda* samples revealed admixture from *P. napi* clusters in the southern range border.

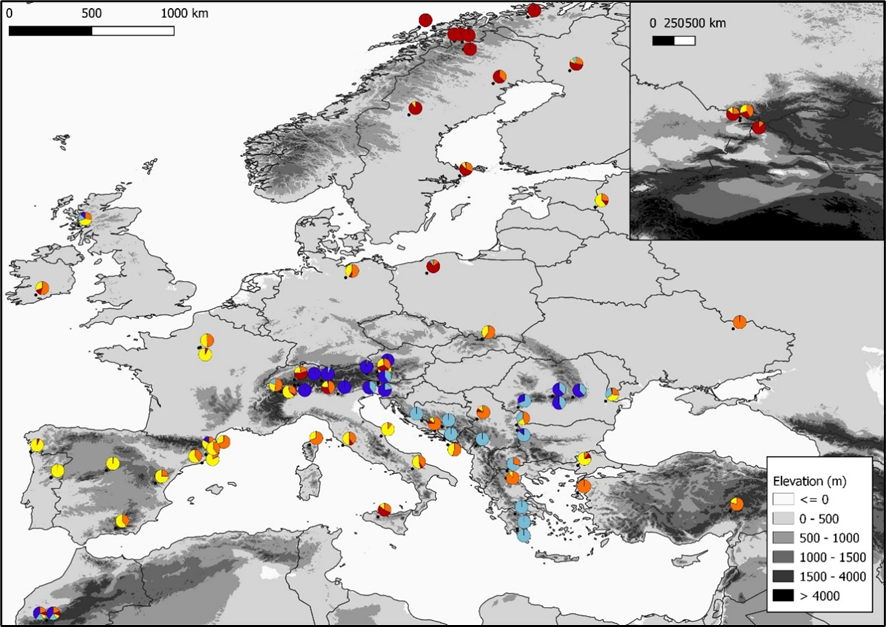
